# Rice Pan-genome Array (RPGA): an efficient genotyping solution for pan-genome-based accelerated crop improvement in rice

**DOI:** 10.1101/2022.01.19.476884

**Authors:** Anurag Daware, Ankit Malik, Rishi Srivastava, Durdam Das, Ranjith K. Ellur, Ashok K. Singh, Akhilesh K. Tyagi, Swarup K. Parida

## Abstract

The advent of the pan-genome era has unraveled previously unknown genetic variation existing within diverse crop plants including rice. This untapped genetic variation is believed to account for a major portion of phenotypic variation existing in crop plants and might be responsible for missing heritability. However, the use of conventional single reference-guided genotyping often fails to capture large portion of this genetic variation leading to a reference bias. This makes it difficult to identify and utilize novel population/cultivar-specific genes for crop improvement. To overcome this challenge, we developed a rice pan-genome genotyping array (RPGA) includes 80K genome-wide SNPs which provides simple, user-friendly and cost-effective solution for rapid pan-genome-based genotyping in rice. The GWAS conducted using RPGA-SNP genotyping data of a rice diversity panel detected total of 42 loci, including previously known as well as novel genomic loci regulating grain size/weight traits in rice. Eight of the identified trait-associated loci (dispensable loci) could not be detected with conventional single reference genome-based GWAS and found to be missing from the commonly used Nipponbare reference genome. WD repeat-containing PROTEIN 12 gene underlying one of such dispensable locus on chromosome 7 (*qLWR7*) along with few other non-dispensable loci was subsequently detected using high-resolution QTL mapping confirming authenticity of RPGA-led GWAS. This demonstrates the potential of RPGA-based genotyping to overcome reference bias. Besides GWAS, the application of RPGA-based genotyping for natural allelic diversity and population structure analysis, seed purity and hybridity testing, ultra-high-density genetic map construction and chromosome level genome assembly, and marker-assisted foreground/background selection was successfully demonstrated. Based on these salient outcomes, a web application (http://www.rpgaweb.com) was also developed to provide easy to use platform for imputation of RPGA-based genotyping data using 3K Rice Reference Panel and subsequent GWAS in order to drive genetic improvement of rice.

## INTRODUCTION

Conventionally, a single reference genome sequence per species is considered enough to understand the complete genetic blueprint of a species. However, availability of multiple genomes per species has revealed presence of extensive sequence diversity predominantly in the form of structural variation (SVs), within individuals of the same species. To account for intra-species variation, the concept of pan-genome was proposed (Tettelin et al., 2005). Pan-genome refers to a complete set of genes of a biological clade (for example a species), which includes both core genes (set of genes present in all individuals of a species) and dispensable genes (set of genes which are individual specific or not present in all individuals of a species). Pan-genomes for major crop species including rice, soybean, wheat, tomato, maize, and *Brassica* are now available and all these pan-genomes contains large number of previously unidentified sequence variants and dispensable genes (Hirsch et al., 2014; Montenegro et al., 2017; Sun et al., 2017; Stein et al., 2018; Zhao et al., 2018; Gao et al., 2019; Alonge et al., 2020; Walkowiak et al., 2020).

The functionally characterized dispensable genes so far have been found to perform a diverse range of crucial functions including tolerance to nutrient deficiency, submergence tolerance, organ size regulation, resistance against pathogens (Xu et al., 2006; Ashikawa et al., 2008; Fukuoka et al., 2009; Hattori et al., 2009; Gamuyao et al., 2012; Maron et al., 2013; Wang et al., 2015). Further, compared to core genes, dispensable genes evolve faster (i.e. higher mutation density and higher synonymous to non-synonymous mutation ratio) indicating the vital role of dispensable genes in providing essential diversity for a species to adapt to diverse environmental conditions. Thus, dispensable genes are believed to be a major contributor to phenotypic diversity and adaptive evolution in crop plants. This makes dispensable genes important targets for crop improvement. Some of the dispensable genes like *Sub1A, Pstol*, etc., have already been utilized in rice breeding programs and have become widely popular owing to the enormous impact of these genes on improving overall crop productivity (Xu et al., 2006; Gamuyao et al., 2012). However, the vast majority of dispensable genes remain un-annotated, making it difficult to decipher the role of these genes in regulating important traits in crop plants. The limited knowledge about the function of dispensable genes could be ascribed to sole reliance on a single reference genome for all kinds of genomic mapping studies especially genome-wide association study (GWAS) and quantitative trait loci (QTL) mapping as it leads to reference bias (Coletta et al., 2021). Leveraging pan-genome-based genotyping for genetic mapping studies can potentailly overcome the reference bias imposed by the use of a single reference genome. However, reserachers still relies on comparison of high-quality *de novo* genome assemblies for high-throughput genotyping of SVs, which makes genotyping large number of samples prohebitvely expensive. Thus there is need to decouple discovery and genotyping phase to perform cost-efficent pan-genome-based genotyping. The recent developemnt of pan-genome graphs provide one such alternative. However, these methods still require high-depth whole genome sequencing and also require extensive computational resources making them unsuitable for most plant-breeding applications.

In the recent past, high-density SNP genotyping arrays have became widely popular for large-scale genotyping in crop plants due to simple procedure, fast-turnaround time, simple data-analysis and limited requirment of high-performance computing clusters (HPCCs) (Bianco et al., 2016; McCouch et al., 2016; Li et al., 2019). Despite these advantages, SNP arrays developed so far in most crop plants including rice, assays variants identified using a single reference genome and, therefore, can not capture most of the dispensable gene variation existing in rice germplasm (Chen et al., 2014; Singh et al., 2015; McCouch et al., 2016; Schmidt et al., 2017; Thomson et al., 2017; Torkamaneh et al., 2019). This limits the utility of high-density SNP arrays for large-scale pan-genome-based genotyping. Thus, a pan-genome-based SNP-array that can tag genetic variation (SNPs/InDels/SVs) from both core as well as the dispensable genome, will be a crucial step forward toward the practical utilization of pan-genomes in crop improvement.

Keeping the aforementioned scenario in mind, here we outline the development of “Rice Pan-genome Genotyping Array” (RPGA), a novel pan-genome-based SNP genotyping array that can efficiently capture haplotype variation from the entire 3K rice pan-genome representing diverse population (*indica,* tropical/temperate *japonica, aus* and aromatic, etc.). We further demonstrate its application for sucessfully overcoming reference bias in high-resolution GWAS and QTL mapping to delineate novel genomic loci modulating traits of agronomic importance (for instance, grain size/weight) in rice. Finally, a user-friendly web portal “Rice Pan-genome Genotyping Array Analysis Portal” (RAP) was also developed to provide researchers with easy to use interface for performing imputation and conducting pan-genome-based GWAS using genotype data generated with RPGA. Thus, RPGA and RAP together provide an end-to-end genotyping solution for accelerated genomics-assisted breeding and crop improvement in rice.

## RESULTS

### Designing of Rice Pan-genome Genotyping Array (RPGA)

A rice pan-genome array (80K) design is based on 3K rice pan-genome, which includes the complete Nipponbare genome (IRGSP 1.0/MSU release 7: 373 Mb) and 12 pseudo-chromosomes containing genomic sequence specific to different sub-groups of 4 rice subpopulations (*indica*, *japonica*, *aus* and aromatic) as well as admixed accessions (Sun et al., 2017). RPGA assays total of 80504 SNPs including 60026 SNPs from 12 Nipponbare chromosomes and 20478 SNPs from 12 pseudo-chromosomes of 3K rice pan-genome. The 80504 RPGA SNPs display even distribution throughout the rice pan-genome with ∼5 SNPs in every 100 kb genomic interval. In the case of the Nipponbare genome, chromosome 1 being the longest, harbors the highest number of SNPs (7348) whereas, chromosome 9 being the shortest harbors the least (4704) number of SNPs (**Figure 1a, b**). About 25% (22066) and 47% (37960) of the said 80504 RPGA-SNPs are from genic and intergenic regions of the Nipponbare reference genome, respectively (**Figure 1c**). The remaining 28% (20478) of SNPs belong to the twelve pseudo-chromosomes (dispensable/population-specific genomic sequences) from the 3K rice pan-genome (**Figure 1c**).

**Figure 1.**
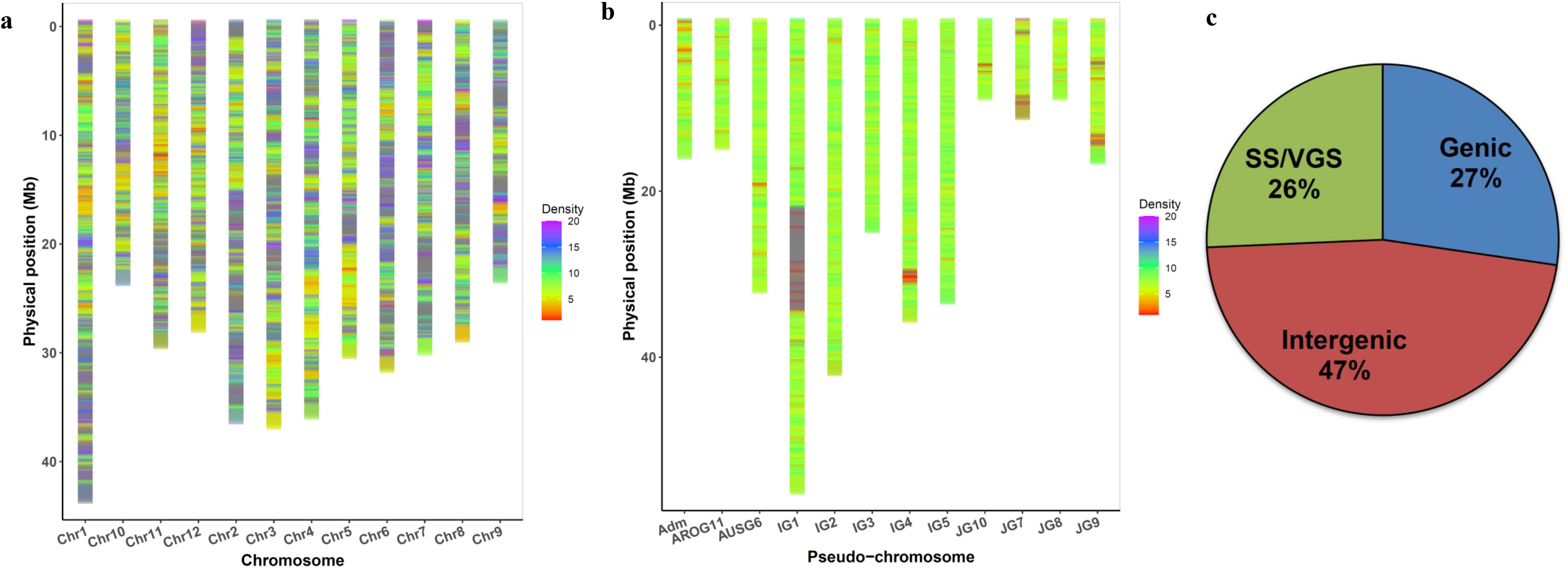
Pan-genome-wide distribution and structural annotation of SNPs tiled on rice pan-genome array. **a)** The density of SNPs in 100 kb bins across twelve chromosomes from Nipponbare reference genome. **b)** The density of SNPs in 100 kb bins across sub-population/varietal group-specific contigs of 3K rice pan-genome. **c)** Proportionate distribution of SNPs across genic and intergenic regions of the Nipponbare genome as well as across sub-population/varietal group-specific sequences (SS/VGS) identified in 3K pan-genome. (Adm: Admixed Group; AROG: Aromatic Group; AUSG: *Aus* group; IG: *Indica* Group; JG: *Japonica* Group)

### Large-scale validation of RPGA

The large-scale validation of RPGA was performed using 188 RILs (Sonasal × Pusa Basmati 1121 F_12_ mapping population) and a diversity panel consisting of 275 accessions representing cultivated and wild Indian rice accessions. In the case of both RILs and diversity panel, very high proportion of SNPs i.e. ∼ 57% (46523) and ∼ 60% (48133) of total 80504 RPGA-SNPsrespectively, were classified as Poly High Resolution (PHR). In addition to PHR SNPs, about 15% (12618) and ∼7% of SNPs (5746) were classified as Off Target Variants (OTVs) in the diversity panel and RILs, respectively, which were then processed using the OTV caller provided as part of Axiom Analysis Suite to obtain genotype calls (presence/absence) (**Figure 2**). Thus, total of 60751 (75.46%) and 52269 (65.68%) of total 80504 SNPs from the diversity panel (275 accessions) and RILs (188 individuals) genotype datasets, respectively, were selected finally for the downstream analysis (**Supplementary Table 1, 2**). More than 98% concordance between RPGA- and whole genome resequencing-based SNP genotyping data of four accessions. Further, > 99% concordance was detected for technical replicates genotyped with RPGA, indicates high-reproducibility of RPGA with extremely low genotyping error call rate and thus, can be reliably used for large-scale genotyping in rice. The selected RPGA-SNPs found to have high minor allele frequencies (MAFs > 0.15) suggesting their highly informative nature (**Supplementary Figure 1a**). The design of RPGA and its genotyping potential across natural and mapping population was optimized for capturing maximum level of subpopulation-specific variations and thus expected to provide high-degree of polymorphism irrespective of germplasm (varietal group and sub-populations) used. The SNP markers belonging to pseudo-chromosomes of different sub-populations/varietal groups displayed high polymorphic potential as reflected by their MAFs (**Supplementary Figure 1b**). This suggests a highly informative nature RPGA for genotyping of diverse rice accessions belonging to different rice subpopulations. Further details regarding the large-scale validation potential of RPGA including optimization of RPGA-SNP genotyping call success rate, and reproducibility/informativeness of valid RPGA-SNP genotyping data across accessions/RILs are provided as **Supplementary Results.**

**Figure 2.**
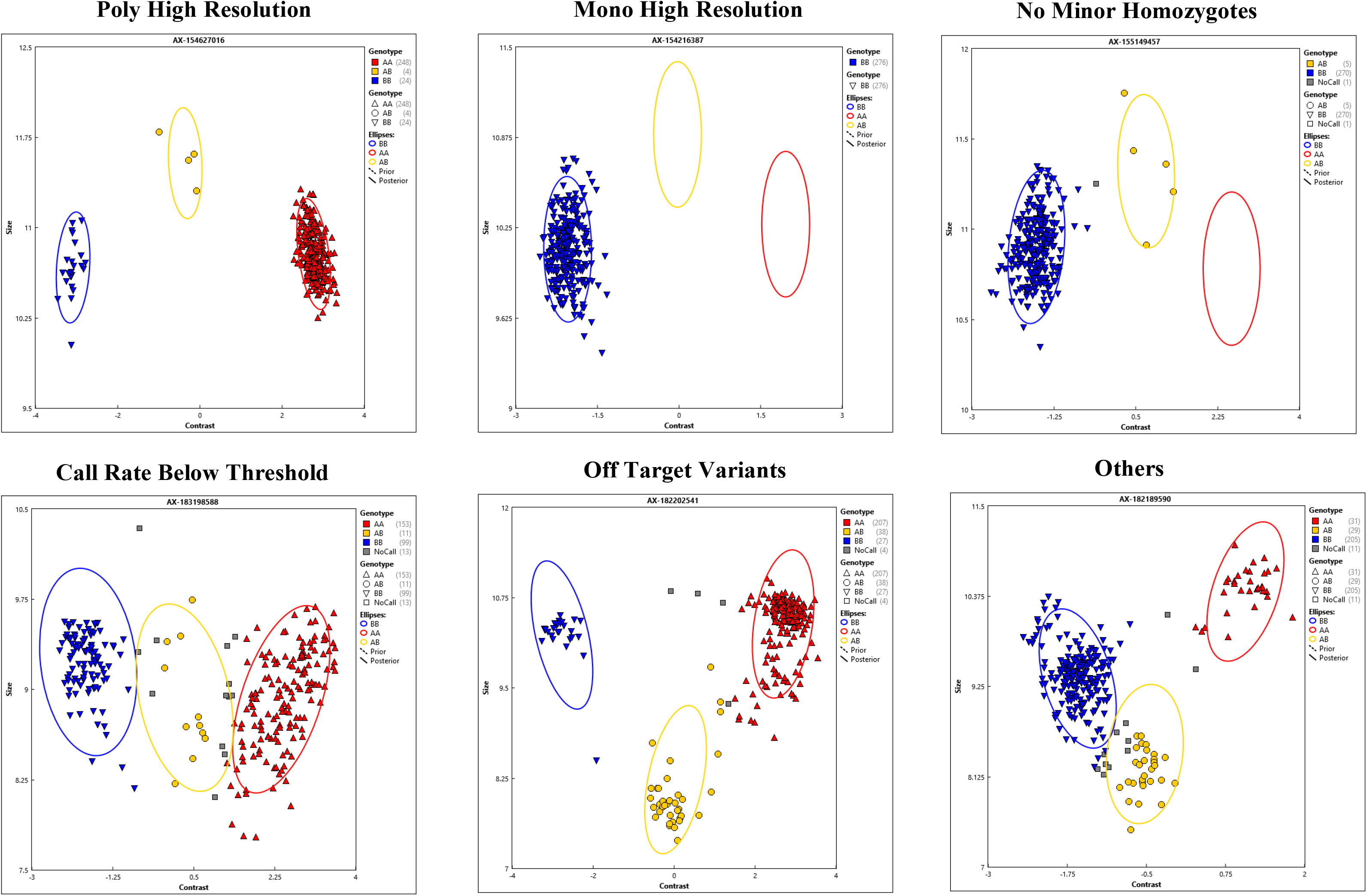
Representative images of six different SNP classes identified using diversity panel genotyping data generated with rice pan-genome genotyping array (RPGA).

### Genetic mapping of sub-population specific (dispensable) sequences from 3K rice pan-genome to twelve rice chromosomes

To develop a framework map for 3K rice pan-genome, an ultra-high-density genetic linkage map using the aforesaid RPGA-based SNP genotyping data of 188 RILs (Sonasal × PB 1121 F_12_) was generated. The RIL genotype contained a total of 46523 PHR SNPs evenly distributed throughout the 12 Nipponbare chromosomes as well as in 12 subpopulation-specific pseudo-chromosomes from 3K rice pan-genome. These SNPs provide uniform genomic coverage of the entire rice pan-genome with ∼7 SNPs/100 kb. Further, filtering for missingness, segregation distortion, the error-rate and duplicate marker was performed to obtain 13792 unique SNPs across 12 Nipponbare chromosomes and 12 sub-population specific pseudo-chromosomes. Initially, only 9100 SNPs belonging to 12 Nipponbare chromosomes were utilized for genetic linkage map construction. This yielded a genetic linkage map spanning a total genetic distance of 1416.4 cM with an average inter-maker distance of 0.1 cM (**Supplementary Table 3**). Chromosomes 1 (LG1) and 7 (LG7) was found to be the longest and shortest spanning 179.4 and 82.6 cM, respectively.

Further, the genetic map was re-constructed using 13793 SNPs, which includes 9100 SNPs from 12 Nipponbare chromosomes as well as 4693 SNPs from 12 pseudo-chromosomes. The genetic linkage map spanned a total genetic distance of 2312.2 cM with an average inter-maker distance of 0.2 cM reflecting the ultra-high-density nature of the constructed genetic map (**Figure 3; Table 4**). Chromosomes 1 (LG1) and 7 (LG7) was found to be the longest and shortest spanning 311.5 and 137.6 cM, respectively (**Supplementary Table 4**). This ultra-high-density genetic map generated using markers from the entire 3K rice pan-genome enabled the identification of genetic positions of 4693 SNPs belonging to subpopulation/pseudo-chromosomes relative to the SNPs from the Nipponbare reference genome. These 4693 SNPs tagged > 93% contigs from 12 sub-population specific pseudo-chromosomes, harboring thousands of important dispensable genes. Therefore, RPGA-based ultra-high-density genetic linkage map provides a good estimate of genetic locations of these contigs from 12 sub-population specific pseudo-chromosomes.

**Figure 3.**
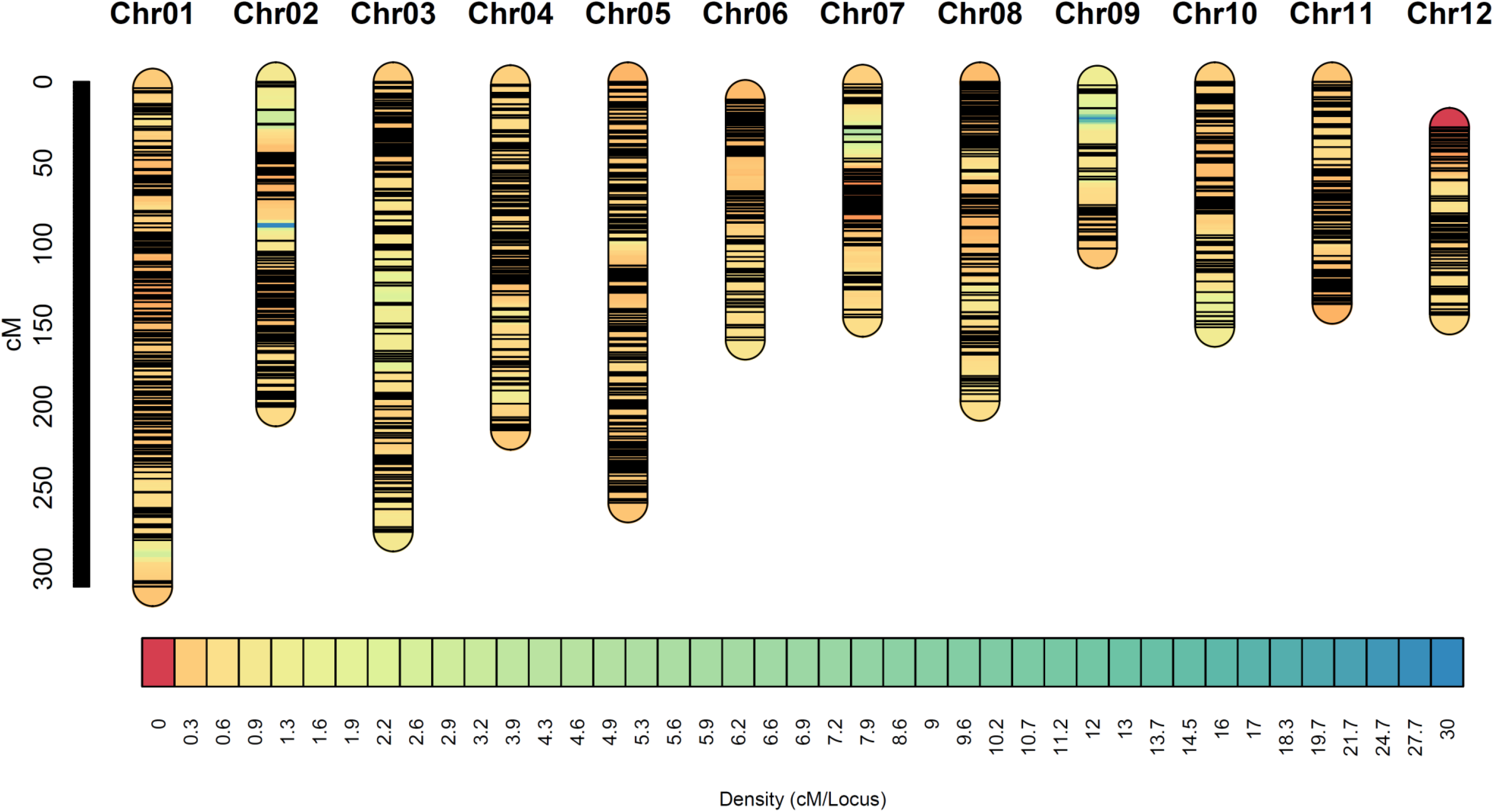
Ultra high-density genetic linkage map of a RIL mapping population (Sonasal × Pusa Basmati 1121) developed using genotype data generated with rice pan-genome genotyping array (RPGA). SNPs are represented as black horizontal lines on each linkage group. The vertical scale at the right depicts the genetic distance in cM. The color scale at the bottom represents the genetic distance represented by each locus across linkage groups in the form of density (cM/Locus).

Further, approximate physical locations of 4693 SNPs from 12 sub-population specific pseudo-chromosomes relative to Nipponbare reference genome were determined using an integrated approach. This involves BLAST search against six genome assemblies, namely Nipponbare (*japonica*), Nagina 22 (*aus*), IR 64 (*indica*), Sonasal, Basmati 334 and Dom Sufid (aromatic), pair-wise LD estimates and genetic positions of SNPs derived from aforementioned high-density genetic linkage map. The information on the physical positions of contigs and dispensable genes within these contigs is vital for a diverse range of genomic applications. This is especially important in both GWAS and QTL mapping studies, to identify dispensable novel genes (unable to detect from Nipponbare reference genome) underlying QTLs governing important agronomic traits.

### Validation of RPGA-based ultra-high-density genetic map using *de novo* genome assembly of “Sonasal”

To assess accuracy of RPGA-based Sonsal × PB 1121 ultra-high-density genetic map, the *de novo* whole genome assembly of one of the parental accession i.e., “Sonasal” was generated using Oxford Nanopore long-read sequencing. The Sonasal genome assembly (304 contigs) spanning 368.2 Mb with 41020 annotated genes. The Sonasal genome assembly revealed high contiguity with N50 of 4.02 Mb and 96.5% recovery of Benchmarking Universal Single-Copy Orthologs (BUSCO). These BUSCO statistics are comparable to previously assembled rice genomes including *japonica* cultivar Nipponbare (98.4%), *indica* cultivar R498 (98.0%), Basmati cultivar Basmati 334 (97%) and *Sadri* cultivar Dom Sufid (97%) (Choi et al., 2020). About 93% (∼ 343 Mb) of assembled 368.2 Mb contig sequences of 12 chromosomes of Sonasal genome was efficiently anchored onto 12 linkage groups of an ultra-high-density genetic linkage map. Further, whole-genome sequence alignment revealed high degree of microsynteny between Sonasal and diverse rice genomes including Nipponbare. This ascertains the high-accuracy and usefulness of RPGA-based ultra-high density genetic linkage map for anchoring *de novo* assembled contigs (**Figure 4**). The comparison of high-quality *de novo* Sonasal contigs with RPGA-based ultra-high-density genetic linkage map revealed high concordance in marker order with > 99.8% markers agreeing with each other. This confirmed the high-quality of RPGA-based ultra-high density genetic linkage map for their immense use in developing the high-quality chromosome-level genome assemblies as well as identification/mapping of sub-population specific dispensable novel sequences (genes) from 3K rice pan-genome on chromosomes of rice.

**Figure 4.**
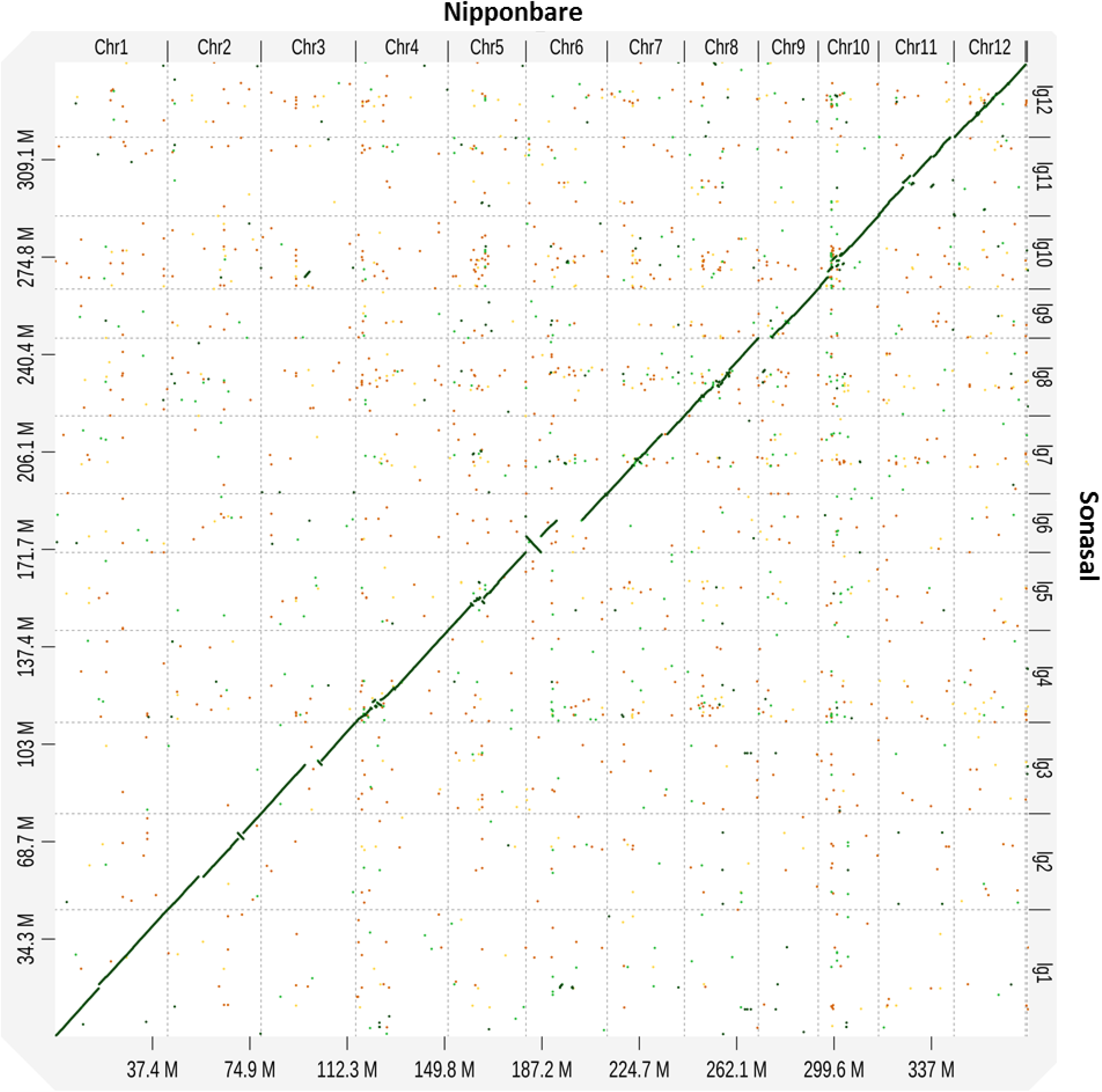
Dot plot illustrating syntenic relationship between the Sonasal (aromatic) draft genome and Nipponbare (*japonica*) reference genome. The 12 LGs (linkage groups) correspond to 12 Sonasal chromosomes.

### Assessment of natural allelic diversity, evolutionary pattern and population structure in a diversity panel

The principal component analysis (PCA) was conducted using the RPGA-SNP genotyping data of 271 rice accessions alone as well as along with 3045 accessions from 3K rice genome (Wang *et al*., 2018b). The first two principal components (PCs) explained > 90% of genetic variation. The first PC separated *indica* distinctly from *japonica* and aromatic/traditional Basmati accessions, whereas the second PC separated *aus* and *indica* accessions (**Figure 5a, b**). The majority of these rice accessions belonged to *indica*, *aus* and aromatic subpopulations, whereas only few are represented from the *japonica* subpopulation. Similar to previous reports, *indica* accessions were found to be genetically closer to *aus*. Interestingly, both Indian aromatic*/*traditional Basmati and evolved Basmati accessions clustered distinctly from aromatic/Basmati accessions present in the 3K rice reference panel. The Indian aromatic*/*traditional Basmati accessions were found genetically closer with *japonica* and *aus* accessions. Whereas the evolved Basmati were found to cluster between *indica* and aromatic*/*traditional Basmati accessions, following their evolution from cross-hybridization between traditional Basmati varieties with superior grain qualities and *indica* varieties with dwarf height, early flowering and high yield (Singh et al., 2018).

**Figure 5.**
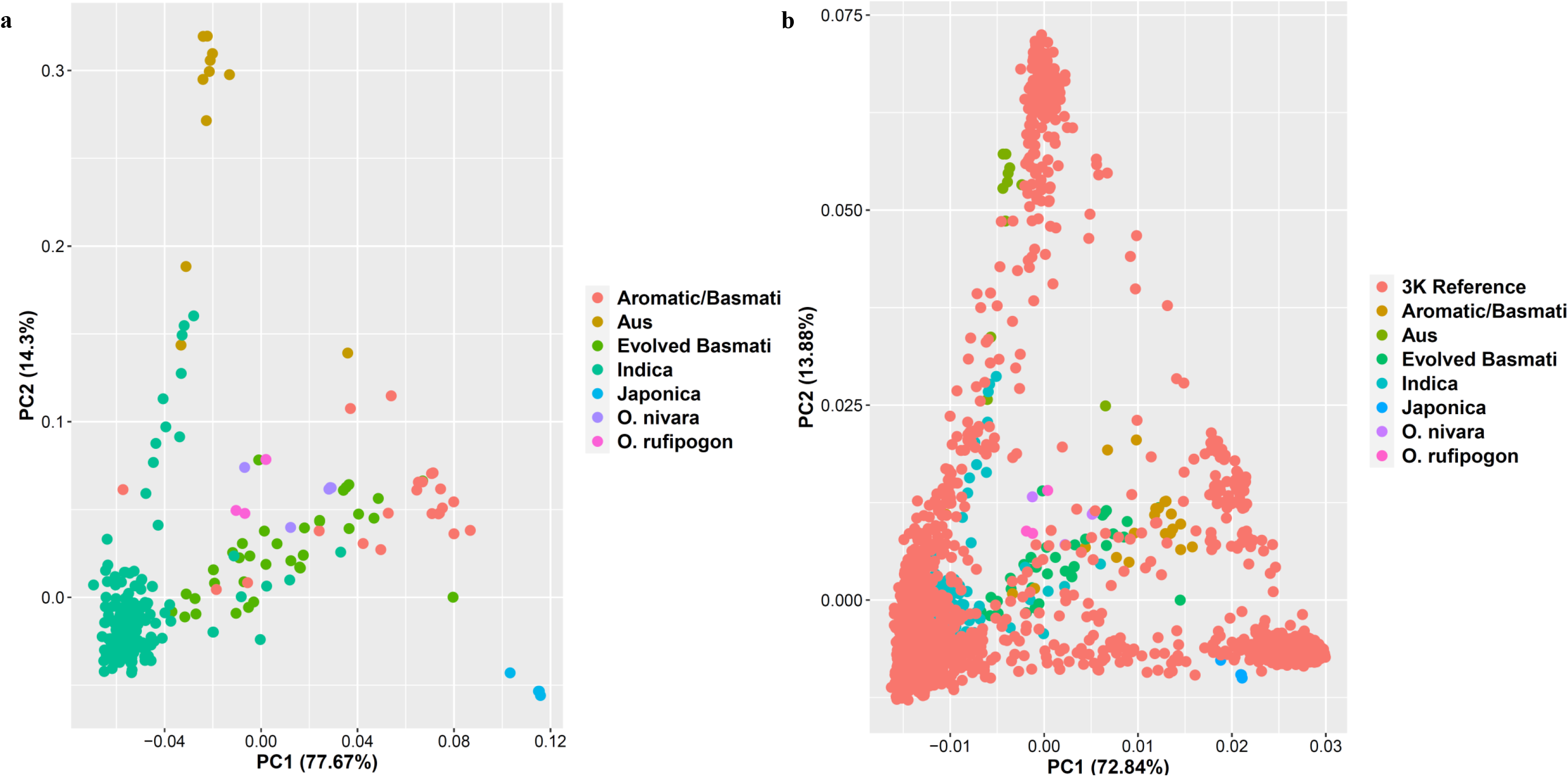
Principal component analysis (PCA)-based molecular diversity analysis of a diversity panel with rice accessions representing different rice sub-populations (*indica, japonica, aus,* and aromatic*/*Basmati). **a)** PCA of 271 rice accessions, where, principal component 1 (PC1) and PC2 explains 77.67% and 14.43% genetic variation, respectively, in the accessions analyzed. **b)** PCA of 271 Indian rice accessions along with 3K rice genome reference panel, where, PC1 and PC2 explains 72.84% and 13.88% genetic variation, respectively, in the accessions analyzed.

Subsequently, the RPGA-SNP genotyping data generated for 271 rice accessions (60751 SNPs) was further analyzed using fastSTRUCTURE to unravel the fine population structure among rice accessions under study. The results revealed the existence of five distinct population groups in rice accessions. These population groups corresponded to two *indica* sub-groups (*INDI* and *INDII*), and one population group each of *aus*, aromatic/traditional Basmati (ARO*/*TR-BAS) and evolved Basmati (EV-BAS), respectively. Based on ancestry cutoff of 65%, accessions were classified into one of the five distinct population groups (*INDI*, *INDII*, *AUS*, ARO*/*TR-BAS and EV-BAS) and admixture classes, admixed *Indica* (*INDI - INDII*), admixed *Indica-aus*, and admixed *Indica-*aromatic (**Figure 6a**). The two *indica* subpopulation groups corresponded to *Xian*/*Indica-2* (*XI*-2) and *XI*-3 from South Asia and Southeast Asia, respectively, which are previously reported along with two other *indica* subpopulation groups (*XI-1A* from East Asia and*XI-1B* constitutes modern varieties of diverse origins) (Wang et al., 2018b). Further, the *indica* rice germplasm accessions especially from India belong to either of the two aforesaid *indica* subpopulations or evolved as a result of cross-hybridization within these subpopulations or with any of the remaining subpopulations. This suggests the indigenous nature of the most widely cultivated Indian *indica* rice accessions. Apart from this, traditional Basmati accessions and aromatic landraces from north-eastern India were found to cluster together as a group close to traditional Basmati distinct from modern evolved Basmati/aromatic accessions. Contrary to this, evolved Basmati/aromatic accessions were closely related to the *IND1* subpopulation, confirming their origin from cross-hybridization between *IND1* and traditional Basmati accessions. The neighbor-joining (NJ) phylogenetic tree generated also confirmed the existence of five distinct population groups apart from three admixed populations (**Figure 6b**).

**Figure 6.**
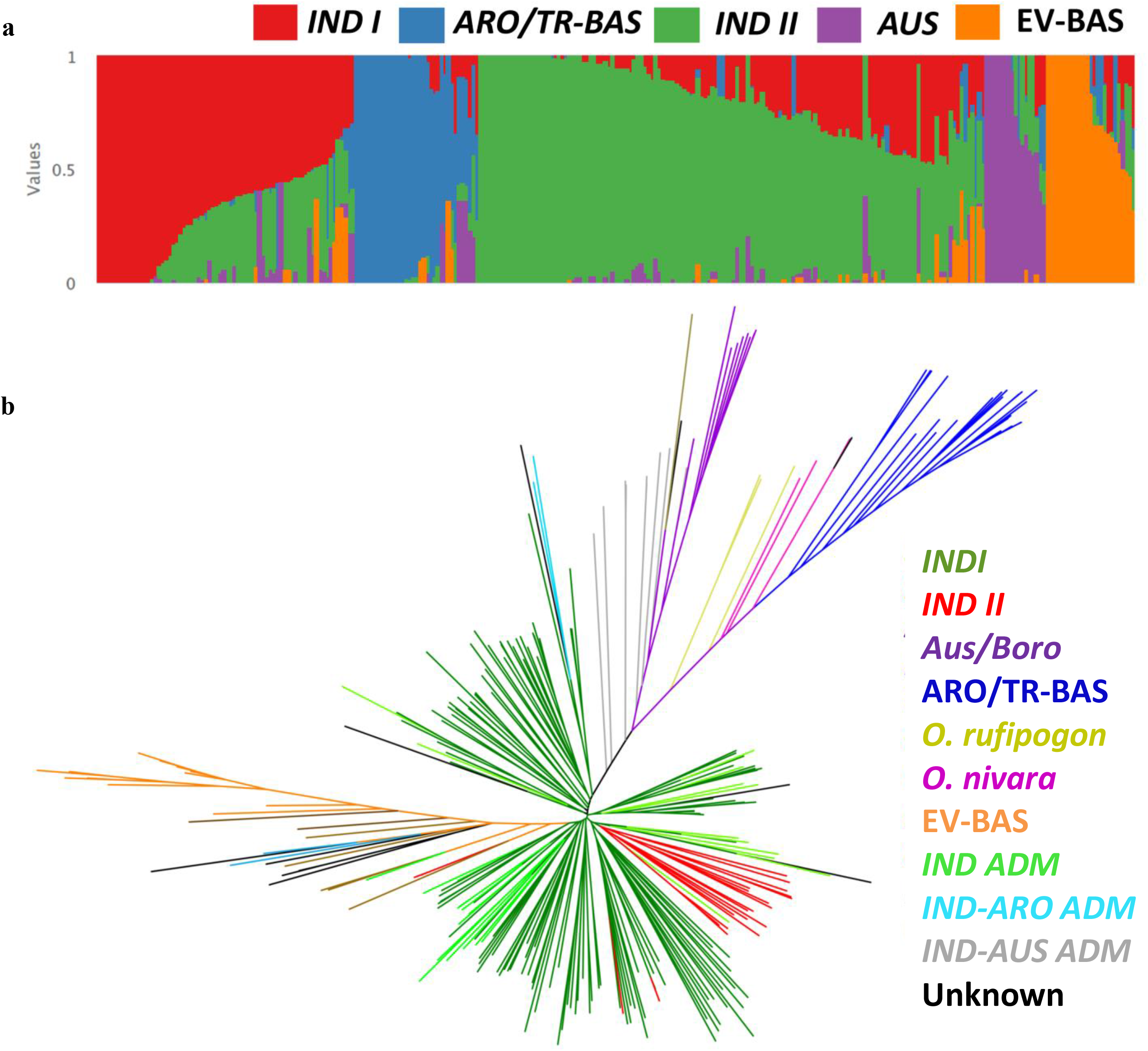
Population structure and neighbor-joining tree-based molecular diversity in a diversity panel of Indian rice accessions. **a**) Population structure of 265 diverse rice accessions with K=5, Inferred populations are color-coded with five different colors. **b**) Neighbor-joining tree of 271 diverse rice accessions belonging to three different cultivated and wild rice species viz. *O. sativa, O. nivara* and *O. rufipogon*. The branches of phylogenetic are colored to depict their respective genetic ancestry. ADM: Admixture, ARO: Aromatic, *Aus/Boro*: *Aus/Boro* ecotypes, EV-BAS: Evolved Basmati, *IND*: *Indica,* TR-BAS: Traditional Basmati.

### The utility of RPGA for conducting pan-genome-based GWAS

To establish the utility of RPGA for conducting pan-genome-based GWAS, the analysis was performed using the RPGA-SNP genotyping data of 203 germplasm accessions (selected out of the aforesaid 275 rice accessions), phenotyped for four different grain size/weight traits, i.e. grain length, grain width, length-to-width ratio and thousand-grain weight. These 203 accessions displayed a wide phenotypic variation for all four grain size/weight traits measured. The grain length of accessions ranged from 4.4 to 8.3 mm with a mean ± S.D of 5.9 ± 0.8 mm, whereas, grain width varied from 1.3 to 2.9 mm with a mean ± S.D of 1.9 ± 0.3 mm. The thousand-grain weight of accessions ranged from 11.6 to 41.3 g with a mean ± S.D of 23.1 ± 4.5 g (**Supplementary Figure 2, 3**). Pan-genome**-**based GWAS was performed using the RPGA genotyping data of 63002 SNPs that includes 48256 SNPs from the Nipponbare reference genome and 14746 SNPs from the subpopulation-specific pseudo-chromosomes, across 206 rice accessions. The pan-genome-based GWAS detected a number of significant associations for all four studied grain size/weight traits. These associations belong to Nipponbare chromosomes as well as sub-population-specific pseudo-chromosomes, which together constitute the 3K rice pan-genome.

The pan-genome-based GWAS detected many previously known gene loci regulating grain size and weight of rice. These include major grain size gene *GRAIN SIZE 3* (*GS3*) which is known to regulate grain size i.e., grain length, grain width and length-to-width ratio (Fan et al., 2009; Takano-Kai et al., 2009; Lu et al., 2013). Apart from *GS3, POSITIVE REGULATOR OF GRAIN LENGTH 1 (PGL1)*, a known positive regulator of grain length in rice is also found to be associated with grain length (Heang and Sassa, 2012a,b). Similarly, previously known major grain size/weight genes, *GW5* and *OsPPKL1* were also found strongly associated with both grain width as well as thousand-grain weight (Weng et al., 2008; Gao et al. 2019). These results not only reaffirmed an important role of previously known grain size/weight genes such as *GS3*, *GW5* and *OsPPKL1* genes in regulating grain size and grain weight in rice accessions but also validated the authenticity of RPGA-based marker-trait association study (**Figure 7**; **Supplementary Figure 4, 5, 6, 7; Supplementary Table 5, 6, 7, 8, 9**).

**Figure 7.**
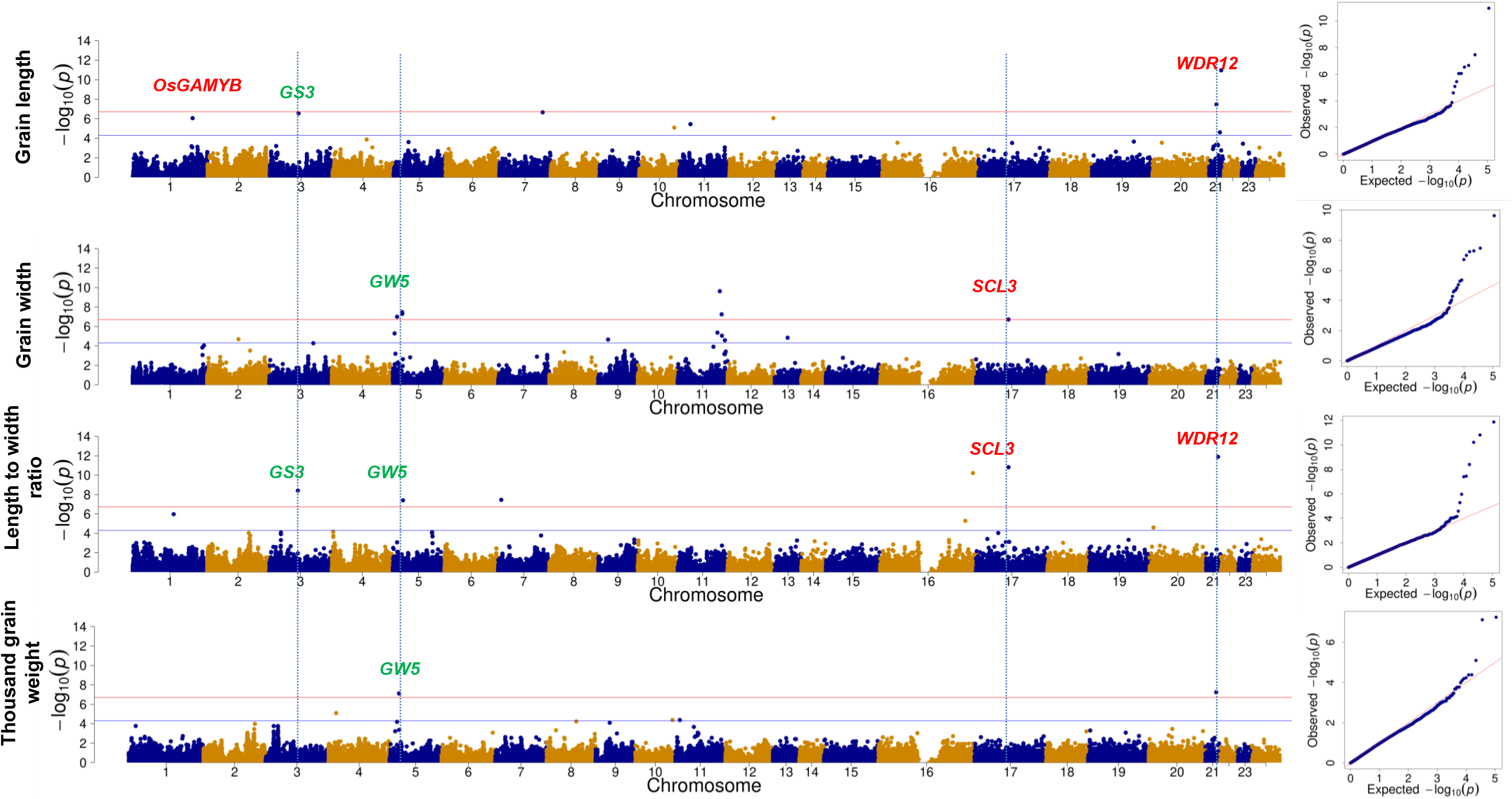
Manhattan plots and QQ plots depicting results of genome-wide association study (GWAS) for grain length, grain width, length-to-width ratio, thousand-grain weight trait. Chromosomes 1 to 12 represent twelve Nipponbare chromosomes, whereas, chromosomes 13 to 24 represent twelve sub-population group specific pseudo-chromosomes (Chr13:Adm, Chr14:AROG11, Chr15:AUSG6, Chr16:IG1, Chr17:IG2, Chr18:IG3, Chr19:IG4, Chr20:IG5, Chr21:JG10, Chr22:JG7, Chr23:JG8, Chr24:JG9). Trait associated loci coinciding with previously known grain weight genes are marked in green. The blue horizontal line represents a stringent Benjamini-Hochberg threshold whereas the red horizontal line represents a less stringent threshold. (Adm: Admixed Group; AROG: Aromatic Group; AUSG: *Aus* group; IG: *Indica* Group; JG: *Japonica* Group).

In addition to the aforementioned known genes, many novel genomic loci including 10 loci for grain length, 5 loci for grain width, 3 loci for length-to-width ratio and 9 loci for thousand-grain weight, were detected from the Nipponbare reference genome (**Figure 7; Supplementary Figure 4, 5, 6, 7; Supplementary Table 5, 6, 7, 8, 9**). These include transcription regulators like *OsGAMYB1* (*LOC_Os01g59660*) and *OsRUB1* (*LOC_Os10g11260*) genes involved in ubiquitin proteasome pathway as well as the serine/threonine-protein kinase gene (*LOC_Os11g40970*) modulating grain size/weight in rice (**Details in Supplementary Results**).

### RPGA-based GWAS successfully uncovers novel sub-population specific (dispensable) genes regulating grain size/weight in rice

Apart from loci detected from the 12 Nipponbare chromosomes, multiple SNPs/contigs of pseudo-chromosomes were found to be significantly associated with one or more of the 4 target grain size/weight traits that are repeatedly detected with different statistical models of GWAS. These include 2 loci each for grain length, grain width and grain length-to-width ratio, and 3 loci for thousand-grain weight in rice. Since these SNPs/contigs are known to be partially or completely missing from the Nipponabre reference genome, the physical positions of these contigs were first determined using an integrated genomic approach which leverages information from RPGA-based genetic linkage map, alignment of sub-population specific contigs to multiple reference genome assemblies and pair-wise LD across rice pan-genome (Detailed strategy as per Materials and methods). The candidate genes underlying these loci associated with grain size/weight were subsequently identified. Here three of these vital trait associations for rice grain size/weight are discussed in further details.

### *GW5* associated with rice thousand grain weight

The prominent locus detected from the pseudo-chromosome IG5 (MSP-unaln_IG5∼14354843) was found to be strongly associated with thousand-grain weight. Further, physical location of this locus was identified using the aforesaid integrated approach. Based on this analysis, the physical position of target locus coincides with the 1212 bp InDel (SV) located upstream of *GW5* (calmodulin binding protein-encoding gene) that has been identified previously as a causal variation responsible for regulating grain size/weight (Liu et al., 2017). This further proves the efficacy of the RPGA-based GWAS approach to efficiently identify SVs associated with important agronomic traits in rice.

### *WDR12* associated with rice grain length and grain length-to-width **ratio**

Apart from this, a SNP from *japonica* group 10 (JG10) pseudo-chromosome (MSP-unaln_JG10∼6950498) was repeatedly detected for grain length as well as grain length-to-width ratio (**Figure 7; Supplementary Figure 4, 5, 6, 7; Supplementary Table 5, 6, 7, 8, 9**). The location of SNP unaln_JG10∼6950498 was detected on chromosome 7 with high confidence. Further, to identify underlying candidate genes for this QTL, 5 kb flanking sequences from the location of SNPs were retrieved from genome assemblies of seven different rice accessions belonging to different rice subpopulations. This includes *japonica* (Nipponbare), *indica* (IR 64 and R 498), *aus* (Nagina 22) and aromatic (Sonasal, Basmati 334 and Dom Sufid) subpopulations. The extracted sequences were then subjected to multiple sequence alignment along with the full contig sequences harboring the lead SNPs to determine any variation present within this region, between the aforementioned seven rice accessions. The SNP-unaln_JG10∼6950498 spanned the upstream/promoter regions of two consecutive genes encoding WD repeat-containing PROTEIN 12 (*LOC_Os07g40930*) and X8 domain-containing PLASMODESMATA CALLOSE-BINDING PROTEIN (*LOC_Os07g40940*). A closer look at multiple sequence alignment revealed the presence of multiple InDels within 1.5 kb upstream region of *LOC_Os07g40930*. These InDels are likely to impact the promoter of *LOC_Os07g40930* possibly modulating its expression. Further, the InterPro scan detected multiple WD40 repeats confirming *LOC_Os07g40930* as a member of the WD-repeat WDR12/Ytm1 family. Interestingly, PESCADILLO (PES), a conserved nucleolar protein involved in ribosome biogenesis, is known to perform its function by interacting with BLOCK OF PROLIFERATION 1 (BOP1) and WD REPEAT DOMAIN 12 (*WDR12*) to form the PeBoW (PES-BOP1-WDR12) complex. The reduction in PeBoW proteins was found to suppress cell proliferation and cell expansion in *Arabidopsis* underling its vital role in the regulation of cell cycle for growth and development in plants (Cho et al., 2013; Ahn et al., 2016). Apart from this, members of the WD40 repeat superfamily are known F-box proteins, a structural component of SKP1/Cullin/F-box (SCF) E3 ubiquitin ligase complex. These include *Arabidopsis STERILE APETALA* (*SAP*) gene and its ortholog from cucumber *LITTLELEAF (LL)*. Both of these genes encode WD40 repeat domain-containing F-box proteins and regulate organ size further emphasizing the vital role of WD40 repeat domain-containing proteins in plant growth and development (Wang et al., 2016c; Yang et al., 2018). This evidence indicates the probable role of *LOC_Os07g40930* in cell number/cell size regulation in rice and, therefore, *LOC_Os07g40930* seems to be a probable candidate gene regulating grain length in rice.

### GRAS family transcription factor associated with grain length

For grain length, another important locus was detected on the pseudo-chromosome JG10 (MSP-unaln_JG10∼4245875) (**Figure 7; Supplementary Figure 4, 5, 6, 7; Supplementary Table 5, 6, 7, 8, 9**). However, the contig harboring SNP, unaln_JG10∼4245875 (2.19 kb) was found to be completely missing from the Nipponbare (*japonica*) reference genome but present on the chromosome 6 in genomes of multiple other rice accessions including *indica* (IR 64 and R 498), *aus* (Nagina 22) and aromatic (Sonasal, Basmati 334, and Dom Sufid). The physical position of this SNP locus (QTLs thereof) was further determined with reference to the Nipponbare genome using an integrated genomic approach as described earlier. Similar to SNP-unaln_JG10∼6950498, 5 kb flanking sequences from the location of SNP unaln_JG10∼4245875 in all the aforesaid genomes except Nipponbare (as the sequence is missing from the Nipponbare genome) were retrieved and compared using multiple sequence alignment. The contig harboring SNP-unaln_JG10∼4245875 (2.19 kb) was found to overlap with the GRAS family transcription factor gene (*OsR498G0612839600.01*) in R 498 (*indica*) reference genome. This gene predominantly expresses in leaf, panicle and endosperm of R 498 *indica* rice accession and encodes for 498 amino acid protein. Subsequent, comparison of an *OsR498G0612839600.01* genomic sequence among five rice accessions, revealed 1669 bp insertion in all three aromatic accessions (Sonasal, Basmati 334 and Dom Sufid) compared to both *indica* (IR 64 and R 498) and an *aus* (Nagina 22), leading to truncated protein with 108 amino acids. Interestingly, the GRAS family proteins are previously reported to play a vital role in plant development, including male gametogenesis predominantly by regulating cell division, proliferation and maintenance. One of such GRAS family rice gene, *GRAIN SIZE 6* (*GS6*) */OsGRAS-32/ DLT* is reported to regulate grain width and thousand-grain weight by promoting cell expansion in spikelet hull in rice through modulating both brassinosteroid and gibberellin signaling (Sun et al., 2013). However, a homology study revealed that protein encoded by *OsR498G0612839600.01* is homolog to scarecrow-like protein 3 (*SCL3*) which belongs scarecrow subgroup of the GRAS family. In *Arabidopsis*, SCL3 is known to function downstream of DELA transcription factor (another subgroup of GRAS family) and are known to negatively regulate of gibberellin signaling. *SCL3* plays an important a role in *Arabidopsis* root elongation by regulating cell division in roots (Lee et al., 2016). However, the effect of *SCL3* on seed development is not yet studied in plants. Thus, it would be interesting to study the probable role of *SCL3* in regulating grain size/weight of rice.

Besides, diverse sub-population specific (dispensable) novel candidate genes underlying the grain size/weight loci mapped on pseudo-chromosomes were identified. These include genes encoding multi-domain protein (*LOC_Os10g31770*) for grain width and grain length-to-width ratio as well as dirigent family (*LOC_Os12g12600*) and unknown expressed (*LOC_Os12g12610*) genes for grain weight in rice (**Details in Supplementary Results)**.

### Imputation of RPGA-SNP genotyping data using 3K Rice Reference Panel (RICE-RP)

Imputation of high-quality reference SNP genotyping dataset dramatically increases marker density and has been shown to positively impact GWAS results. As the RPGA is designed to efficiently tag almost all haplotype variation existing in 3K rice panel, the imputation of RPGA-based genotyping data of diversity panel was performed using 3K RICE-RP (Wang et al., 2018a). This increased 48257 to 5231751 SNPs from 12 Nipponbare chromosomes. These SNPs tags > 3500 cloned and functionally characterized genes in rice. Further, 2760746 of these SNPs displayed minor allele frequency (MAF) >0.05, which translates average marker density of ∼ 7.4 SNPs/kb compared to pre-imputation average marker density of ∼ 0.12 SNPs/kb.This result suggests that the SNP genotyping data generated using RPGA can be efficiently imputed to obtain a manifold increase in marker density which can further result in higher resolution and enhanced power of QTL detection in GWAS.

Further, GWAS analysis was repeated for grain length and grain width traits using imputed RPGA genotyping data with a total of ∼2.76 M SNPs to assess the effect of increased marker density on power and resolution of trait association in rice. The post-imputation GWAS for grain length trait detected *Grain Size 3* (*GS3*) locus; however, with a much smaller p-value. Another major locus detected on chromosome 12 also crossed a stringent significance threshold which previously failed to do so (**Supplementary Figure 8**). This indicates the increased power of QTL detection in GWAS conducted using the imputed RPGA-based SNP genotyping data. Interestingly, post-imputation GWAS for grain width trait detected a novel locus on chromosome 8 which remained undetected before imputation (**Supplementary Figure 8**). This highlights the potential of imputed RPGA-based SNP genotyping data to detect novel associations which otherwise would be missed due to low marker-density.

### RPGA-based high-resolution QTL mapping to dissect the genetic basis of grain size/weight variation in aromatic rice

To conduct QTL mapping, 190 mapping individuals and parental accessions selected from a developed F_12_ RIL population (Sonasal × PB 1121) RIL () were phenotyped for grain length, grain width, length-to-width ratio and thousand grain weight. The grain length of RILs ranged from 3.5 to 8.1 mm with a mean ± standard deviation (SD) of 5.4 ± 1.4 mm. The grain width of RILs varied from 1.5 to 2.5 mm with a mean ± SD of 1.9 ± 0.17 mm. Finally, the thousand-grain weight for RILs ranged from 8.8 to 31.3 g with a mean ± SD of 17.2 ± 4.4 g. Further, grain length positively correlated with thousand-grain weight (Pearson’s correlation coefficient, r = 0.74) whereas grain width was found to be negatively correlated with both grain length (r = −0.29) and thousand-grain weight (r = −0.032) (**Supplementary Figure 9, 10**). This suggests thousand-grain weight is predominantly influenced by grain length, whereas, grain width has either little or negligible influence on the thousand-grain weight in the said mapping population.

The previously generated ultra-high-density genetic linkage map (Sonasal × PB 1121 RILs) and high-quality SNP genotyping data (13793 SNPs) were integrated with grain size/weight (grain length, grain width, length-to-width ratio and thousand-grain weight) phenotype data of 190 RILs and parents, to identify potential QTLs and their underlying candidate genes governing said target traits in rice. A total of six QTLs which are genetically mapped on either chromosome 3 or 7 governing four grain size/weight traits were identified **(Figure 8; Supplementary Table 10)**. The phenotypic variation explained (PVE) by these grain size/weight QTLs varied from 13.3 to 42 % **(Supplementary Table 10**). Five of these QTLs showed correspondence with previously known grain size/weight genes. Among these, major grain length QTL, *qGL3* and thousand-grain weight QTL, *qTGW3*, coincides with *GS3* locus (Fan et al., 2006). The grain length QTL*, qGL7* found to harbor previously known grain length gene *GRAIN LENGTH 7* (*GL7*) (Wang et al., 2015; Zhou et al., 2015). Similarly, grain width QTL, *qGW7* contains Os*BZR1* which is a known regulator of grain size in rice (Tong et al., 2009). The QTL detected for length-to-width ratio *qLWR7* harbors *GL3.2,* a previously known grain length regulating gene encoding cytochrome P450 **(Figure 8; Supplementary Table 10)**.

**Figure 8.**
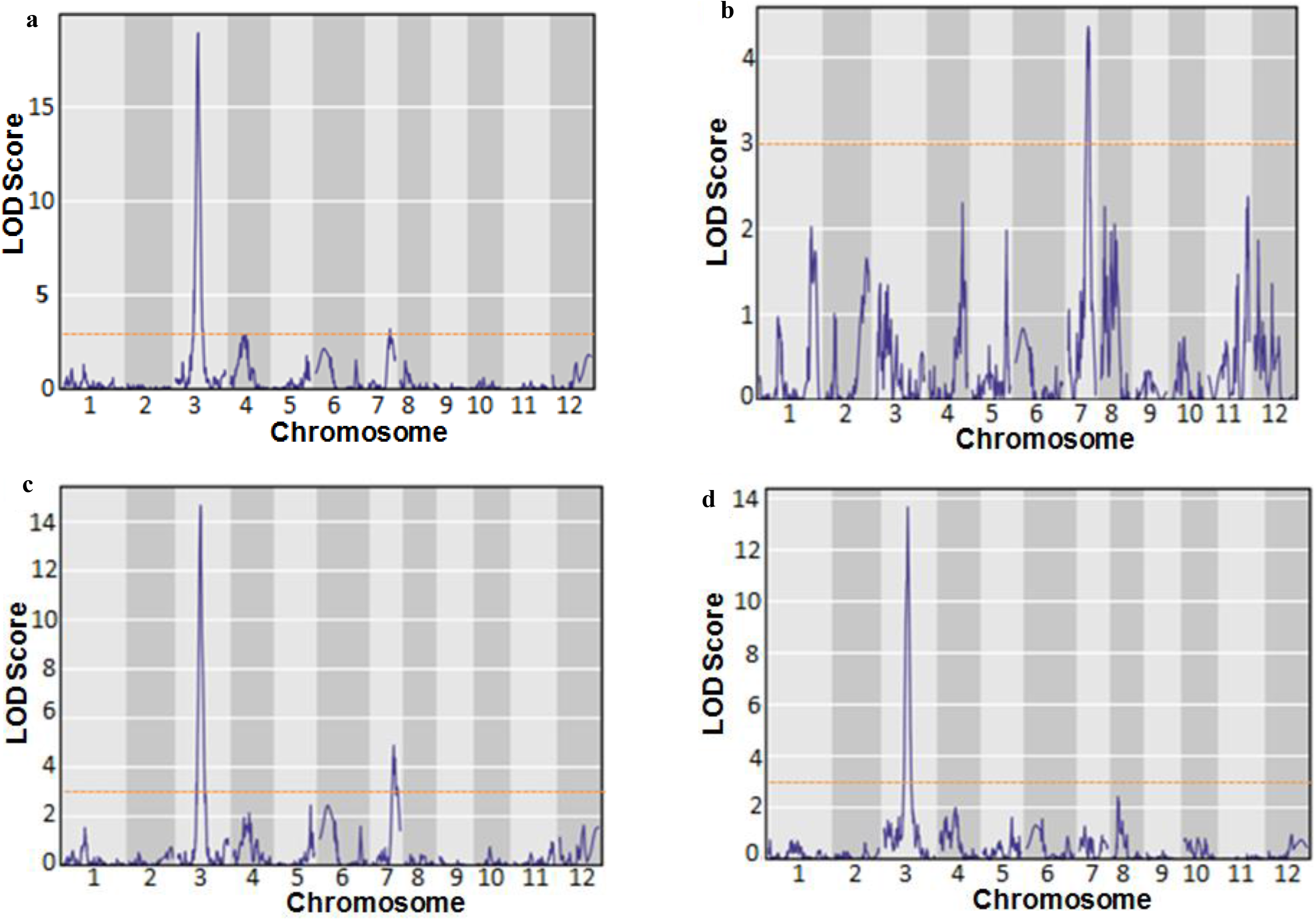
Molecular mapping of QTLs regulating grain size/weight traits in rice. **a)** Thousand-grain weight, **b)** grain length, **c)** grain width, and **d)** length-to-width ratio. The orange color horizontal lines represent the significance threshold calculated using 1000 permutations and red asterisk sign represents the significant QTL peaks.

Interestingly, the only novel QTL identified on chromosome 7 (*qLWR7*) was found to coincide with dispensable locus associated with grain length as well as grain length-to-width ratio as identified in RPGA-based GWAS (**Figure 7; Figure 8c)**. Thus revalidating the authenticity of previously detected dispensable loci using RPGA-based GWAS. As previously discussed, the locus contains WDR12 protein encoding gene (*LOC_Os07g40930*) can be a candidate gene modulating grain size/weight in rice, as WDR12 protein is a vital component of the PeBoW (PES-BOP1-WDR12) complex, which is known to regulate cell proliferation in *Arabidopsis* (Cho et al., 2013; Ahn et al., 2016).

### The utility of RPGA in hybridity testing and genetic background recovery analysis

To demonstrate the utility of RPGA for hybridity testing, multiple F_1_ hybrids generated by cross-hybridization of PB 1121 with different *indica* rice accessions (Vandana, Phule Radha, Pusa 1927, C101A51, and Kalibagh) were genotyped using RPGA. The genome-wide SNP data consisting of 42565 SNP markers were then converted into the Graphical Genotype to visually detect the hybrid lines. Comparison of rice accessions using the parental polymorphic SNP markers across 12 rice chromosomes showed heterozygous nature of analyzed F_1_ lines revealing the usefulness of RPGA in hybridity testing of rice.

Similarly, to demonstrate utility of RPGA for background selection commonly required for marker-assisted breeding and crop improvement of rice, two NILs with Sonasal (recurrent parent) and LGR (donor parent) were genotyped using RPGA-SNP. The RPGA SNP genotyping data of two NILs were then compared with recurrent and donor parental accessions. The graphical genotype for genotyping data of 42565 SNPs in the NILs was made to assess the genomic contribution from both parental accessions. This graphical representation ascertained the advantages of RPGA in higher genetic background recovery analysis, further highlighting the importance of RPGA for marker-assisted crop improvement in rice (**Supplementary Figure 11, 12**).

### Rice Genome Genotyping Array Analysis Portal (RAP) provides a user-friendly interface for analyzing RPGA SNP genotyping data

A web-based application RAP was developed to provide the user with the ability to conduct pan-genome-based GWAS using RPGA-based SNP genotyped data without the need for programming skills. Using RAP, pan-genome-based GWAS can be performed either with existing RPGA genotyping data of diverse rice accessions or user can upload their own preferred RPGA SNP genotyping data. The RAP further enables the user to identify the physical locations of pseudo-chromosome-specific SNPs concerning different available rice reference genomes, which is crucial for the identification of dispensable genes regulating the trait of interest. Further, RAP also enables users to impute RPGA SNP genotyping data using a 3K rice reference panel. The GWAS performed using the imputed SNP genotyping data is especially useful for the identification of novel causal mutations underlying the trait-associated loci in rice. In addition to GWAS, the RAP also enables the user to retrieve SNP genotyping information based on locus IDs of the rice genes (Rice Genome Annotation Project, http://rice.uga.edu/) as well as using the genomic coordinates of 12 chromosomes. Thus, RAP complements the utility of RPGA, and these two altogether provide end-to-end solutions from genotyping to downstream data analysis for their eventual deployment in genomics-assisted crop improvement in rice (**Figure 9**).

**Figure 9.**
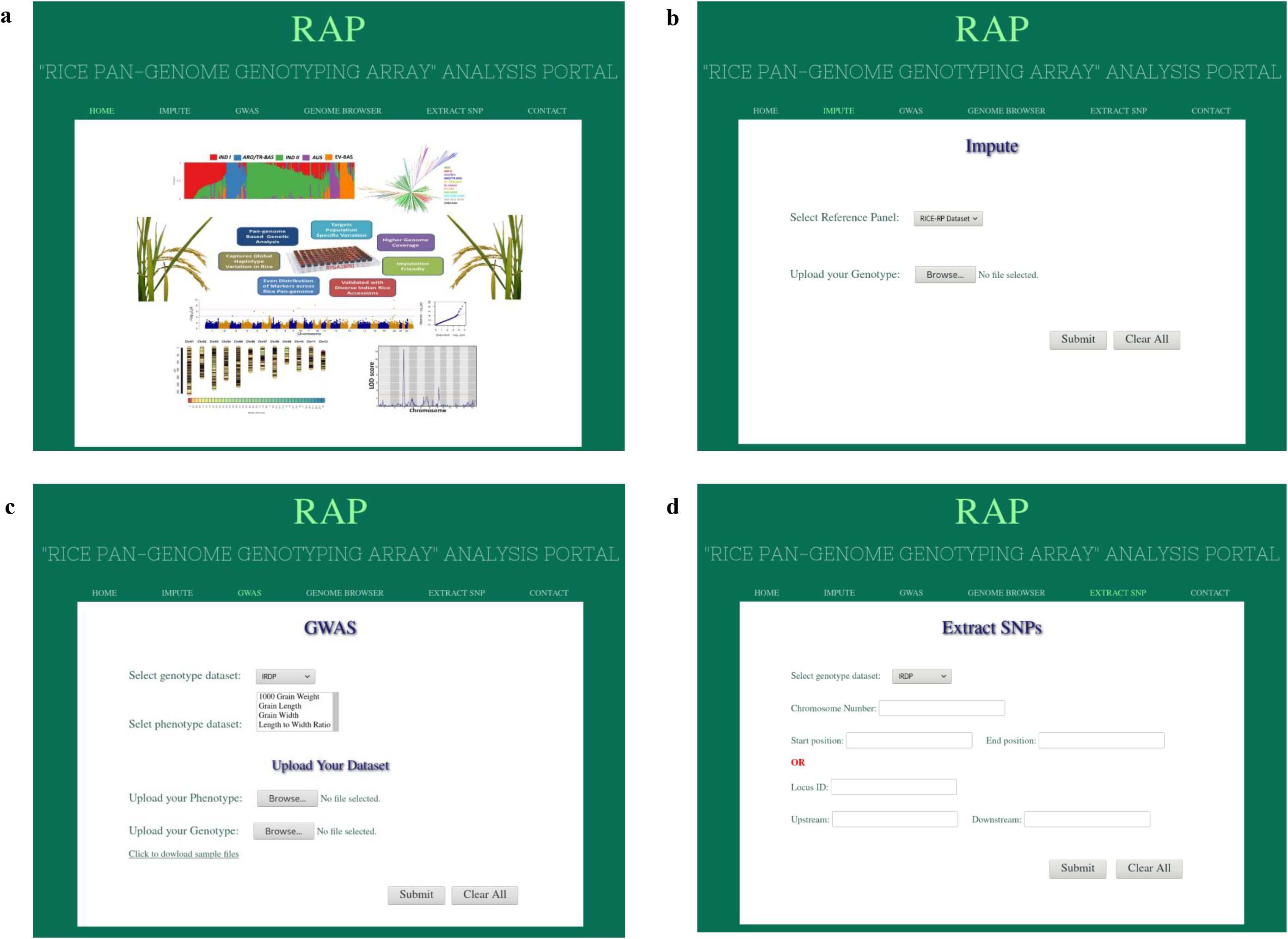
Interface for Rice Pan-genome Genotyping Array Analysis Portal (RAP). **a)** RAP home page, **b)** Impute function available in RAP which enable imputation of user uploaded RPGA genotype data, **c)** GWAS utility available in RAP which enable user to conduct pan-genome-based GWAS, and **d)** RAP enable extraction of SNPs from Locus ID and genomic coordinates.

## DISCUSSION

The recent emergence of pan-genome studies in rice has made evident the presence of extensive dispensable genome diversity in rice (Sun et al., 2017; Wang et al., 2018b; Zhao et al., 2018). Dispensable genes are now known to be the major contributors to adaptive evolution, domestication and overall diversity of agronomically important traits in all major crop plants including rice (Golicz et al., 2016; Tao et al., 2019; Tranchant-Dubreuil et al., 2019). Despite the tremendous potential, leveraging the dispensable gene variation for crop improvement remains a challenging task, especially due to constraints involved in the efficient genotyping of dispensable gene variants in a large number of accessions. Keeping this in mind, we designed “Rice Pan-genome Genotyping Array” (RPGA) to perform rapid, user-friendly and cost-efficient pan-genome-based genotyping in rice. The RPGA is based on a 3K rice pan-genome and, therefore, can efficiently capture variation from both core and dispensable genes, across diverse rice sub-populations (Sun et al., 2017). We demonstrated the utility of RPGA for highly accurate large-scale genotyping of experimental mapping population and diverse natural rice accessions of a diversity panel. The majority of RPGA markers displayed high MAFs suggesting a highly informative nature of RPGA compared to other existing SNP genotyping arrays (Singh et al., 2015; McCouch et al., 2016; Thomson et al., 2017). This suggests suitability of RPGA for diverse high-throughput genotyping applications in rice. Further, a framework map for 3K rice pan-genome generated by integrating RPGA-based ultra-high-density genetic linkage map, a multi-genome alignment of the un-anchored pan-genome contigs and pairwise-LD information between RPGA SNP markers. The importance of such framework map for leveraging crop pan-genomes for multiple genomic-assisted breeding applications have been highlighted previously in maize (Lu et al., 2015). Therefore, the framework-map is expected to be an important step forward toward leveraging 3K rice pan-genome for genomics assisted breeding applications and crop improvement in rice.

Indian rice genepool possesses a large number of highly diverse rice accessions representing different ecogeographical regions across the globe with wide phenotypic variations for multiple economically important agronomic traits. However, comprehensive effort to decipher the natural allelic diversity and population genetic structure among rice accessions was still not completely understood (Kumbhar et al., 2015; Roy et al., 2015; Singh et al., 2016). In this context, RPGA-based SNP genotyping was used to assess the natural allelic diversity and population genetic structure among 271 diverse rice accessions. PCA grouped these accessions into five distinct clusters representing *indica, japonica, aus* and aromatic/Basmati subpopulations which is in accordance with previous studies conducted with global rice accessions (McCouch et al., 2016; Wang et al., 2018b). Further, as expected most of the rice accessions represented from India belonged to either *indica* or *aus* subpopulations, as expected from their morphological characteristics and geographical locations. Interestingly, Indian traditional Basmati accessions were found to cluster distinctly from aromatic rice accessions belonging to both north-eastern India and other parts of the world. Thus, it will be interesting to further identify the precise genes/alleles differentiating Indian Basmati accessions from other aromatic rice accessions. Further, evolved Basmati accessions were found to cluster closer to the *indica* accessions in contrast to traditional Basmati which displayed a closer genetic relationship with *japonica* and *aus* accessions. This can be explained by the breeding history of evolved Basmati varieties, which were developed by cross-hybridization between traditional Basmati varieties with superior grain quality and *indica* varieties possessing favorable traits like dwarf height, early flowering and high yield (Singh et al., 2018). Further, assessment of fine population genetic structure existing within rice accessions revealed the existence of two further sub-groups within *indica* subpopulation i.e. *INDI* and *INDII* corresponding to *Xian*/*Indica-2* (*XI*-*2*) and *XI*-3 from South Asia and Southeast Asia, respectively which are previously reported along with two other *indica* subpopulation groups (*XI-1A* from East Asia, *XI-1B* of modern varieties of diverse origins) (Wang et al., 2018b). This suggests the indigenous nature of the most widely cultivated *indica* rice accessions from India. The evolved Basmati accessions were found to be closely related to the *IND1* subpopulation, confirming their origin from cross-hybridization between *IND1* and traditional Basmati accessions. This suggests the utility of RPGA-based SNP genotyping for efficiently decoding the natural allelic diversity and population genetic structure in order to understand the domestication pattern in rice genepool.

The sole reliance on reference genomes creates a reference bias, which negatively impacts many genomic applications including GWAS (Paten et al., 2017; Coletta et al., 2021). The RPGA-based GWAS conducted in this study detected associations (for diverse grain size/weight traits) from both Nipponbare reference genome and sub-population specific pseudo-chromosomes present in 3K rice pan-genome. The later represents loci partially or completely absent from the Nipponbare reference genome. The previously generated pan-genome framework map allowed us to locate the physical locations of these pseudo-chromosome loci relative to the Nipponbare reference genome and subsequent identification of underlying candidate genes. This demonstrates the potential of RPGA-based GWAS to overcome the reference bias imposed by traditional genotyping based on single reference genome. The RPGA-based GWAS detected many previously known major grain size/weight genes like *GS3* and *PGL1* (grain length and length-to-width ratio) and *GW5* (grain width, length-to-width ratio, and thousand-grain weight) etc., (Fan et al., 2006; Weng et al., 2008; Heang & Sassa, 2012a, 2012b). This not only highlights the major role of these previously known genes in regulating grain size/weight in rice accessions but also validates the ability of pan-genome-based GWAS to detect true associations. The GWAS identified many novel candidate genes including those that are partially or completely absent form Nipponbare genome. These candidate genes similar to previous studies found to be predominantly involved in few major pathways such as ubiquitin-proteasome pathway, brassinosteroid signaling as well as in transcriptional regulation (Li et al. 2018). Subsequently, high-resolution QTL mapping conducted using the RPGA-based ultra-high-density genetic linkage map (Sonasal × PB 1121 RILs) once again identified *GS3* as major locus regulating grain length and thousand grain weight in aromatic rice. Interestingly, a novel chromosome 7 locus (*qLWR7*) regulating length-to-width ratio detected in QTL mapping was also found to overlap with pseudo-chromosome specific QTL detected in RPGA-based GWAS for grain width. This concordance between GWAS and QTL mapping outcomes validates the authenticity of QTLs (from the Nipponbare genome as well as sub-population specific pseudo-chromosomes) identified using RPGA-based GWAS. Thus, RPGA-based GWAS can be efficiently identified using trait-associated genomic loci which otherwise would have been missed using conventional GWAS relying on a single reference genome. The pan genome-based GWAS and QTL mapping overall delineated two known rice grain size/weight genes, namely *GS3* regulating grain length and thousand grain weight and *GW5* governing thousand-grain weight in rice. Besides, multiple promising sub-population specific (dispensable) novel candidate genes modulating grain size/weight for the loci mapped on psudochromosomes were identified. These include *WDR12* (*LOC_Os07g40930*) for grain length and grain length-to-width ratio, multi-domain protein (*LOC_Os10g31770*) for grain width, GRAS family transcription factor (*OsR498G0612839600.*01) for grain length and dirigent family gene (*LOC_Os12g12600*) for grain weight in rice. Among these, the *WDR12* gene underlying the *qLWR7* QTL, validated by both RPGA-based GWAS and QTL mapping, is known to modulate cell number/cell size during growth and development of crop plants and thus appears to be a promising candidate gene regulating grain length and grain length-to-width ratio in rice.

These aforementioned results confirm the utility of RPGA for efficiently leveraging 3K rice pan-genome for diverse genomics-assisted breeding applications including molecular diversity analysis, ultra-high-density genetic linkage map constitution, high-resolution QTL mapping, and GWAS in rice. Further, novel grain size genes/QTLs identified with RPGA-based QTL mapping and GWAS will not only expedite the genetic improvement of Indian rice varieties but also unravel many previously missing links in the genetic and molecular regulation of grain size/weight in rice. Lastly, the “Rice Pan-genome Genotyping Array Analysis Portal” (RAP) developed in this study provides the user with easy to use web interface for handling and analyzing RPGA genotype data. This includes imputation using the 3K Rice Reference Panel and performing pan-genome-based GWAS analysis. This makes the RPGA as an ideal SNP genotyping solution for pan-genome based genetic studies including complex trait dissection and diverse applications in Indian trade and commerce including DNA fingerprinting, genetic purity and hybridity testing. Thus, RPGA and RAP together provide an end-to-end solution for leveraging 3K rice pan-genome for accelerated molecular breeding applications including marker-assisted selection and genomic selection thus enabling genetic enhancement of rice (**Figure 10**).

**Figure 10.**
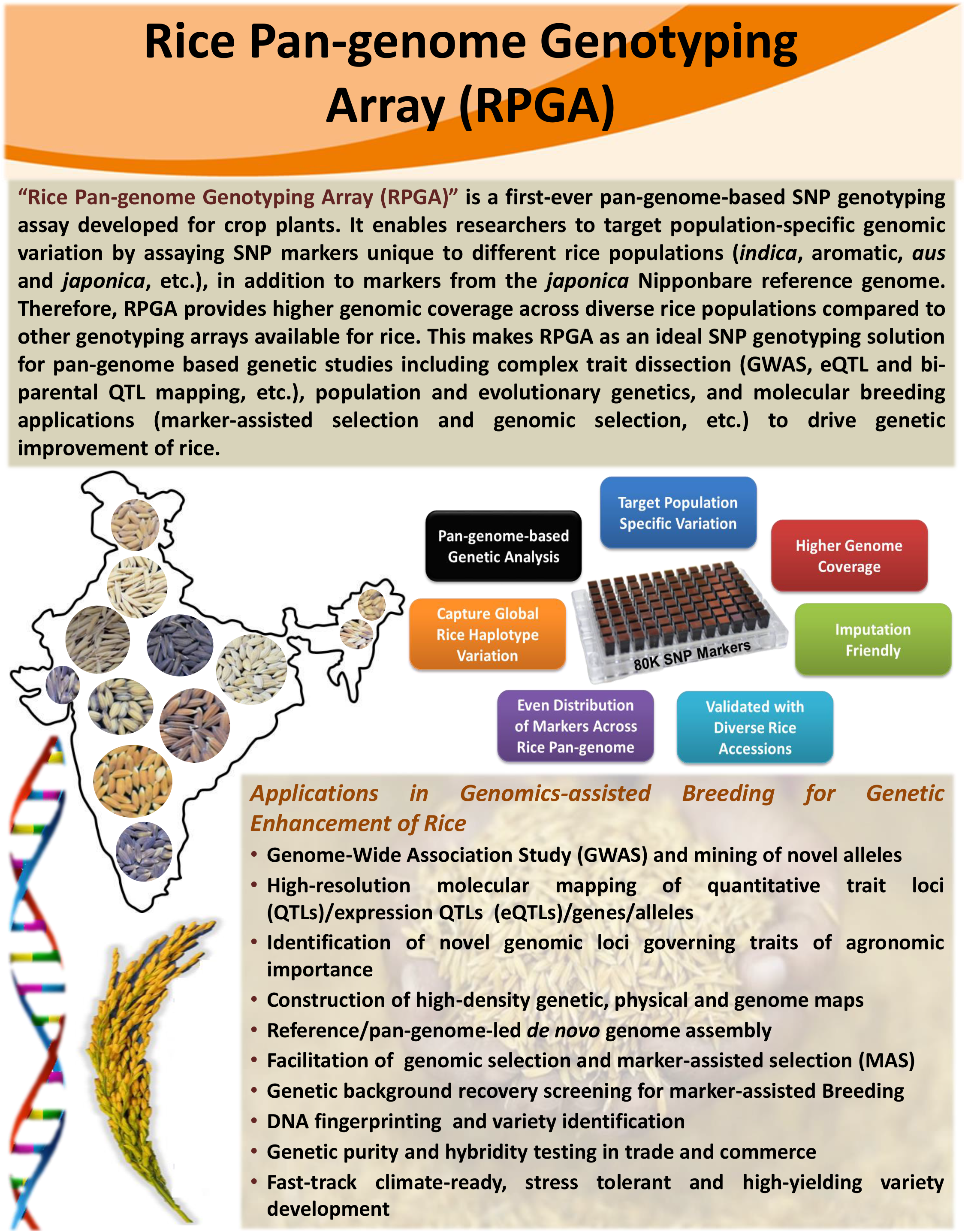
Overview of “Rice Pan-genome Genotyping Array (RPGA)”

## MATERIAL AND METHODS

### Development of a bi-parental mapping population

Pusa Basmati 1121 (PB 1121) and Sonasal, two accessions with contrasting grain size/weight, were utilized for the generation of a bi-parental F_12_ RIL mapping population (190 individuals) varying for grain size and weight traits. PB 1121 is a long-grain Basmati variety with grain length: 9.0 mm, grain width: 2.78 mm and thousand-grain weight: 19.2 g. Sonasal is a short-grain aromatic landrace with grain length: 5.2 mm and grain width: 2.7 mm and thousand-grain weight: 11 g.

### Constitution of the diversity panel

A diversity panel comprising of 271 diverse rice accessions belonging to different rice population (*indica,* tropical/temperate *japonica, aus*, long/short-grain aromatic and wild etc.) was constituted. This panel includes 241 landraces and high-yielding mega-varieties representing different agro-climatic regions of India. In addition, the diversity panel included 24 traditional/ evolved Basmati rice varieties notified by the Government of India and accessions and six accessions of wild rice (three accessions of *Oryza rufipogon* and *Oryza nivara* each). The diversity panel harbors a high level of diversity for a diverse range of traits of agronomic importance including yield (grain size/weight) and a/biotic stress tolerance, etc. Therefore, this diversity panel is ideal for GWAS and genomic selection, especially for grain size/weight trait in rice.

### Phenotypic evaluation

The RIL mapping population (190 individuals) along with parental accessions (PB 1121 and Sonasal) and diversity panel (271 accessions) were grown at the field during the *Kharif* for three consecutive years (2016, 2017 and 2018) in a complete randomized block design with three replications. For the measurement of grain size/dimension traits (grain length and grain width), 50 representative grains were scanned using the Epson Expression 1000XL seed scanner. The mean grain length and mean grain width in millimeters (mm) of each mapping individual/accession were then considered for further analysis. The grain length to grain width ratio of each mapping individual/accession was calculated by dividing mean grain length with mean grain width.

The grain weight (g) was then estimated by measuring the mean weight of 1000 mature dried grains (at 10% moisture content) selected from each accession/individual with three replications. The various statistical parameters, including frequency distribution, coefficient of variation (CV) and broad-sense heritability (H^2^) of grain size/weight traits among accessions/individuals were estimated using SPSSv17.0.

### Designing of a RPGA for SNP genotyping

A rice pan-genome array (80K) was designed to tag haplotype variation from the entire 3K rice pan-genome which includes the complete Nipponbare genome (IRGSP 1.0/MSU release 7: 373 Mbps) and genomic sequence specific to rice pan-genomes representing different sub-groups of 5 rice sub-populations (*indica,* tropical/temperate *japonica, aus* and aromatic, etc.) (Sun et al., 2017) **(Supplementary Figure 13)**. To select SNPs representing the Nipponbare genome, SNPs were retrieved from all the major publicly available rice sequence variation datasets which include: I) 3,000 Rice Genomes Project: ∼ 40 M SNPs from 3243 resequenced global rice accessions (Wang et al., 2018b), II) RiceVarMap: ∼ 6.5 million SNPs from 1479 resequenced Chinese rice accessions (Zhao et al., 2015), and III) High-Density Rice Array (HDRA): ∼ 0.7 M SNPs (700K SNPs) from 1953 genotyped rice accessions (McCouch et al., 2016). Further, to identify SNPs from remaining rice pan-genome sequences (different rice sub-group specific genomic contigs), the publicly available short-read sequence data for 400 Indian rice accessions (NCBI-SRA) including PB 1121 and Sonasal (Kumar et al. 2021) were aligned against the rice 3K pan-genome sequence (Sun et al., 2017) using BWA program (Li and Durbin, 2009). The high-quality SNPs (∼ 0.5 M) were then identified utilizing standard GATK best practices workflow for variant calling (McKenna et al., 2010; Poplin et al., 2017). From this constituted SNP resource, 80504 SNPs distributed uniformly (50 kb genomic intervals) across 3K rice pan-genomes were finally screened following diverse selection criteria and quality filter parameters for tiling on the RPGA **(Supplementary Method)**. The RPGA includes 2000 SNPs from 164 functionally characterized known genes associated with grain size/weight, grain quality and grain aroma traits in rice.

### RPGA genotyping, genotype data analysis and imputation

The genotyping of both RIL population (190 individuals and two parental accessions) and the diversity panel (271 accessions) was performed using the 80K RPGA on GeneTitan^®^ Multi Channel (MC) instrument (Affymetrix, USA) following manufacturer’s instructions (**Supplementary Method**). Subsequent quality control and genotype calling was carried using Axiom Analysis Suite 2.0 (Affymetrix, USA) following Axiom best practices workflow (**Supplementary Method**). The genotype data of the diversity panel was imputed based on rice reference panel (RICE-RP) generated by combining 3K Rice Genome dataset (18M SNPs from 3024 accessions) and HDRA dataset (700K SNPs from 1568 accessions) using IMPUTE2 (Howie et al., 2009) as per Wang et al. (2018a).

### High-density genetic linkage map construction

To construct a high-density genetic linkage map, 80K RPGA-derived genotype data of SNPs showing polymorphism between two parental accessions as well as 190 mapping individuals of a RIL population (Sonasal × PB 1121) were analyzed in the R/qtl program (Broman et al., 2003). The SNP genotyping data of RILs were used to construct a preliminary genetic map with the ordered groupings of marker-pairs in the individual linkage groups at a minimum logarithm of odds (LOD) of 8 and maximum pair-wise recombination fraction (RF) of 0.40 **(Supplementary Method)**. After removing the erroneous SNP genotyping data of RILs and subsequent optimal reordering of markers in the linkage groups, a RPGA-based high-density genetic map was constructed finally.

### *De novo* genome assembly and annotation of a draft genome of Sonasal

For long-read Nanopore and short-read Illumina sequencing, high-quality genomic DNA isolated from the young leaf samples of a short-grain aromatic rice variety Sonasal was used for constructing genomic DNA libraries using the SQK-LSK109 ligation kit (company name, country) and Illumina TruSeq DNA sample prep kit (Illumina, USA), respectively following the manufacturer’s protocol. The libraries prepared were sequenced using the FLO-MIN106 (R9.4) flowcells on PromethION 24 (P24) Nanopore platform and following the paired-end sequencing workflow of Illumina HiSeq2000 platform (Illumina, USA) (**Supplementary Method**).

The high-quality Nanopore long-reads were corrected using Canu (Koren et al., 2014) and further assembled using Flye genome assembler (Kolmogorov et al., 2019), and finally polished with Recon v1.4.11 (https://github.com/isovic/racon) and Medaka (https://github.com/nanoporetech/medaka) for different rounds. BUSCO v4.0.5 (Simão et al., 2015) was used to assess the completeness of draft genome assembly of Sonasal (**Supplementary Method**). The synteny between the assembled Sonasal draft genome and the Nipponbare reference genome (IRGSP 1.0) was analyzed using D-GENIES program (Cabanettes and Klopp, 2018). Subsequently, comprehensive structural annotation of Sonasal draft genomic sequences was performed using MAKER annotation pipeline (Holt and Yandell, 2011), RepeatMasker v4.0.7 (Ref), SNAP (Korf, 2004) and Augustus (Stanke et al., 2008). The high-density genetic linkage maps developed from Sonasal × PB 1121 RIL mapping population were finally used for the scaffolding of annotated contigs with Chromonomer v1.10 (Catchen et al., 2020) (**Supplementary Method**).

### PCA and LD analysis

For PCA, the genotype data of SNPs common to both 271 Indian rice accessions (RPGA dataset) and rice reference panel with 3032 global rice accessions (3000 Rice Genomes Project, 2014) were retrieved and combined. The PCA was performed on combined as well as aforesaid individual SNP genotype dataset with TASSEL v5.0 (Bradbury et al., 2007). The PCA plot was then generated using the ggfortify R package. LD statistics for the RPGA-derived SNP genotype dataset were calculated using PLINK v2.0 (Chang et al., 2015) with a window size of 50 kb. The LD-decay plot was then generated with R version 3.4.1.

### Population structure and molecular diversity analysis

To decipher the population structure in 271 diverse Indian accessions, a core SNP dataset was generated with the LD pruning procedure implemented in PLINK v2.0 (Chang et al., 2015) following Supplemental Methods. The core SNP dataset (5812 SNPs) was analyzed using the variational Bayesian model of fastSTRUCTURE v1.0 with default parameters and number of clusters (K) ==1 to 15. Finally, the replicates of chosen clusters (most-likely K value) once finalized by best chose function of fastSTRUCTURE were summarized with the CLUMPK (http://clumpak.tau.ac.il/).

### GWAS of grain size/weight

GWAS for grain size/weight traits was performed using five different models (CMLM, MLMM, SUPER, FarmCPU and BLINK) implemented in GAPIT v.3 (Wang and Zang 2019). Before performing GWAS, SNP genotype data were filtered to ensure less than 10% missing genotyping data both for accessions and markers. Besides, markers with less than 5% minor allele frequency (MAF) were also eliminated. For all the five methods, the kinship matrix internally calculated in GAPIT was utilized to account for relatedness between accessions. Further, the top two principal components were used as covariates in statistical models to account for the population structure existing within the diversity panel. Additionally, to balance the trade-off between stringency and false negatives two different significance thresholds [a stringent Benjamini-Hochberg threshold (P ≤ 1.5E^-7^) and a less stringent suggestive threshold (P ≤ 5E^-5^)] were utilized. A grain size/weight QTL detected in more than one method with GWAS was considered true positive.

### Identification of candidate genes underlying trait associated genomic loci

For identification of candidate genes underlying grain size/weight associated loci from the Nipponbare reference genome, haplotype blocks associated with the most significant SNPs were identified with PLINK v2.0 (Chang et al., 2015) and all genes within the haplotype blocks considered as potential candidate genes. Further, candidate genes were prioritized based on the presence of major effect mutations within genes as well as the putative function of these genes.

However, candidate genes underlying associated loci from subpopulation-specific pseudo-chromosomes (MSPs either absent or have unknown physical positions in Nipponbare reference genome) were determined using an integrated approach based on multiple rice reference genomes as per the Supplemental Methods and considered further as novel MSPs associated with grain size/weight in rice.

### QTL mapping

QTL mapping was performed by integrating the high-quality SNP genotyping data across 144 RIL mapping individuals (Sonasal × PB 1121) and high-density genetic linkage map with grain size/weight trait phenotype data of respective RILs utilizing r/qtl package (Broman et al., 2003). Genotype probabilities for these markers were then calculated with calc.genoprob() function utilizing the. Further, interval mapping with Kosambi mapping function was conducted utilizing the scanone() function with Haley-Knott Regression, assuming normal phenotypic distribution. The LOD significance threshold was further determined with 1000 permutations for each trait independently. Further, the QTL model refinement was performed for the detection of additional linked QTLs. The QTL model was then finalized by stepwise forward selection and backward elimination to identify best fit the QTL model for given data. The phenotypic variation explained (PVE) and estimated effect sizes of each QTL was determined by performing *ANOVA* on the final selected model. The 95% confidence interval for each QTL peak was then determined as 1.5 LOD units flanking the peak marker (Mangin et al., 1994; Dupuis and Siegmund, 1999).

### Development of RPGA Analysis Portal (RAP)

RAP (http://www.rpgaweb.com) was developed based on Apache HTTP server (version 2.4.6) integrated with PHP (version 7.3.3) and MySQL (version 8.0.15), on a server machine with Centos 8 operating system. HTML and CSS were used to create a responsive front-end interface whereas PHP JavaScript and R were used to develop the backend of RAP. The imputation pipeline previously described by Wang et al. (2018a) was further integrated to conduct efficient imputation of RPGA genotype data based on rice reference panel (RICE-RP) generated by combining 3K Rice Genome dataset (18M SNPs from 3024 accessions) and High-Density Rice Array dataset (700K SNPs from 1568 accessions). In addition, the GAPIT pipeline (Lipka et al., 2012) and JBrowse (Buels et al., 2016) was integrated to conduct GWAS analysis and visualize variants across the rice genome.

## Supporting information

Supplementary Results

## ACKNOWLEDGMENTS

Financial support for this study was provided by a research grant from the Department of Biotechnology (DBT), Government of India (102/IFD/SAN/2161/2013-14). AD acknowledge the DBT, India for research fellowship award.

