## Supplemental Figure 1 for "Rice Pan-genome Array (RPGA): an efficient genotyping solution for pan-genome-based accelerated crop improvement in rice"

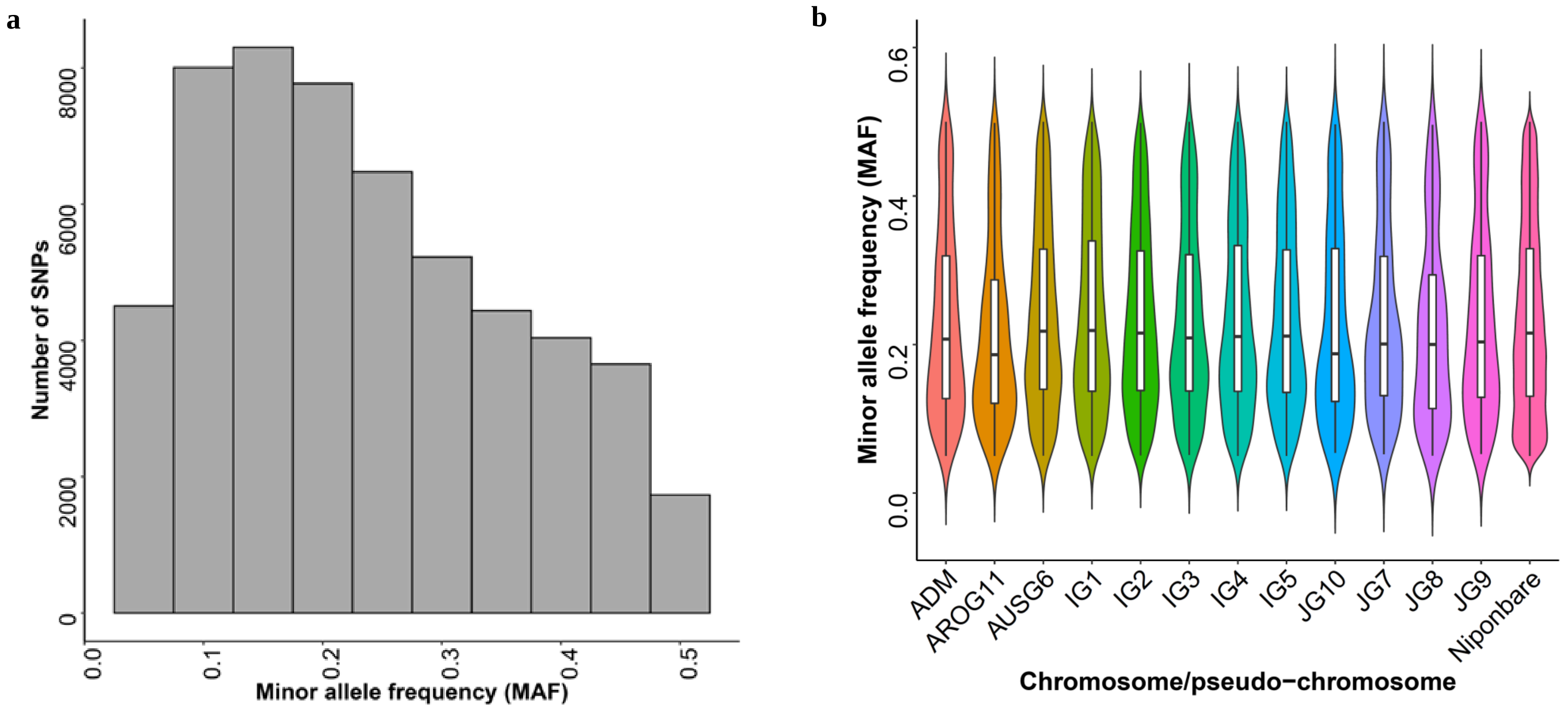


**Supplementary Figure 1. Informativeness of of SNPs genotyped in a diversity panel with rice pan-genome genotyping array (RPGA). a)** Minor allele frequencies (MAFs) of Poly High-Resolution SNPs recorded in a a diversity panel of 271 Indian rice accessions. **b)** Minor allele frequencies (MAFs) of SNPs from Nipponbare reference genome as well as from twelve pseudo-chromosomes from 3K rice pan-genome representing population-specific/unbalanced SNPs. (Adm: Admixed Group; AROG: Aromatic Group; AUSG: *Aus* group; IG: *Indica* Group; JG: *Japonica* Group).
