## Supplemental Figure 2 for "Rice Pan-genome Array (RPGA): an efficient genotyping solution for pan-genome-based accelerated crop improvement in rice"

a

b

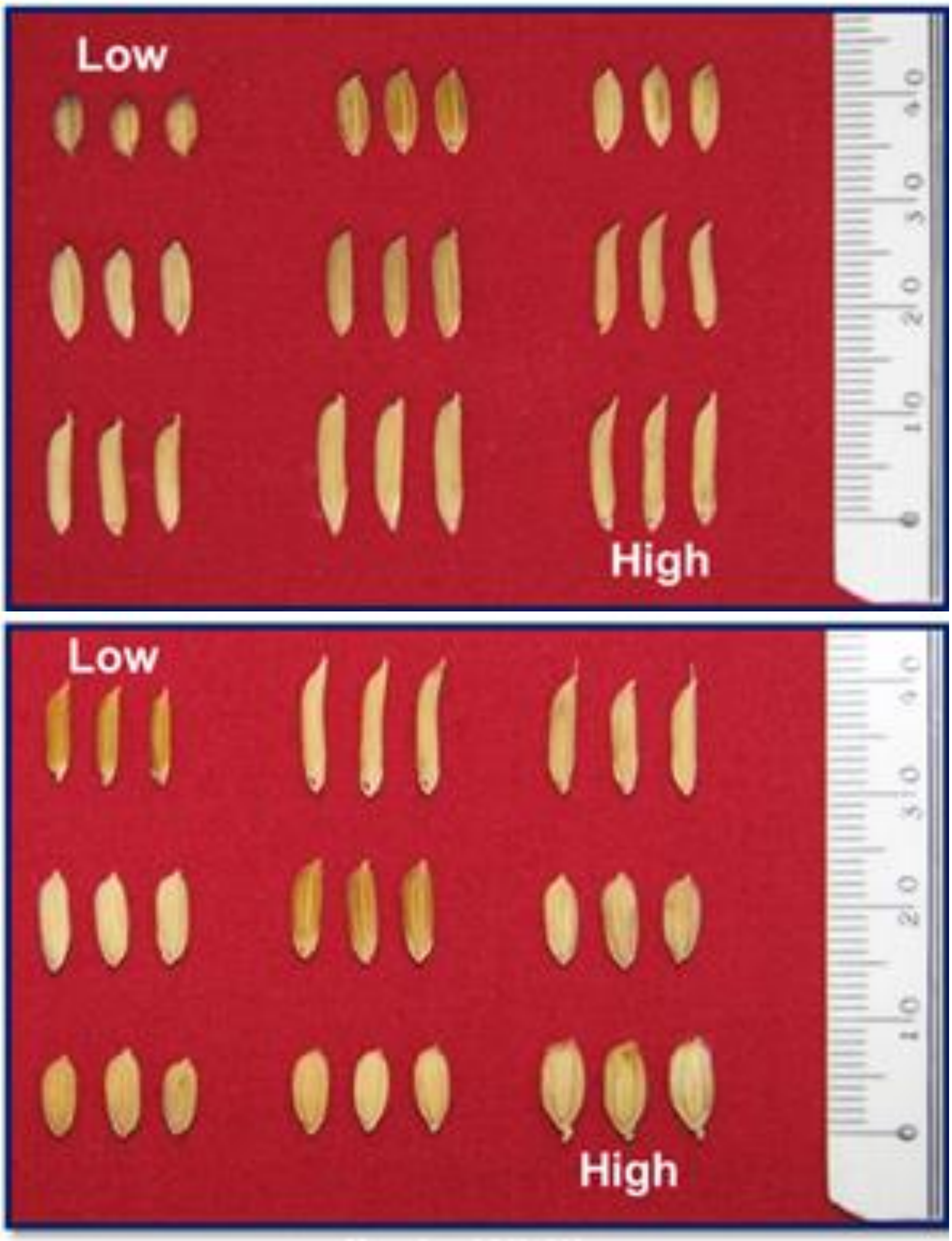


**Supplementary Figure 2. Grains of representative set of rice accessions selected from a diversity panel of 271 Indian rice accessions displaying wide variability for a) grain length and b) grain width traits.**
