## Supplemental Figure 3 for "Rice Pan-genome Array (RPGA): an efficient genotyping solution for pan-genome-based accelerated crop improvement in rice"

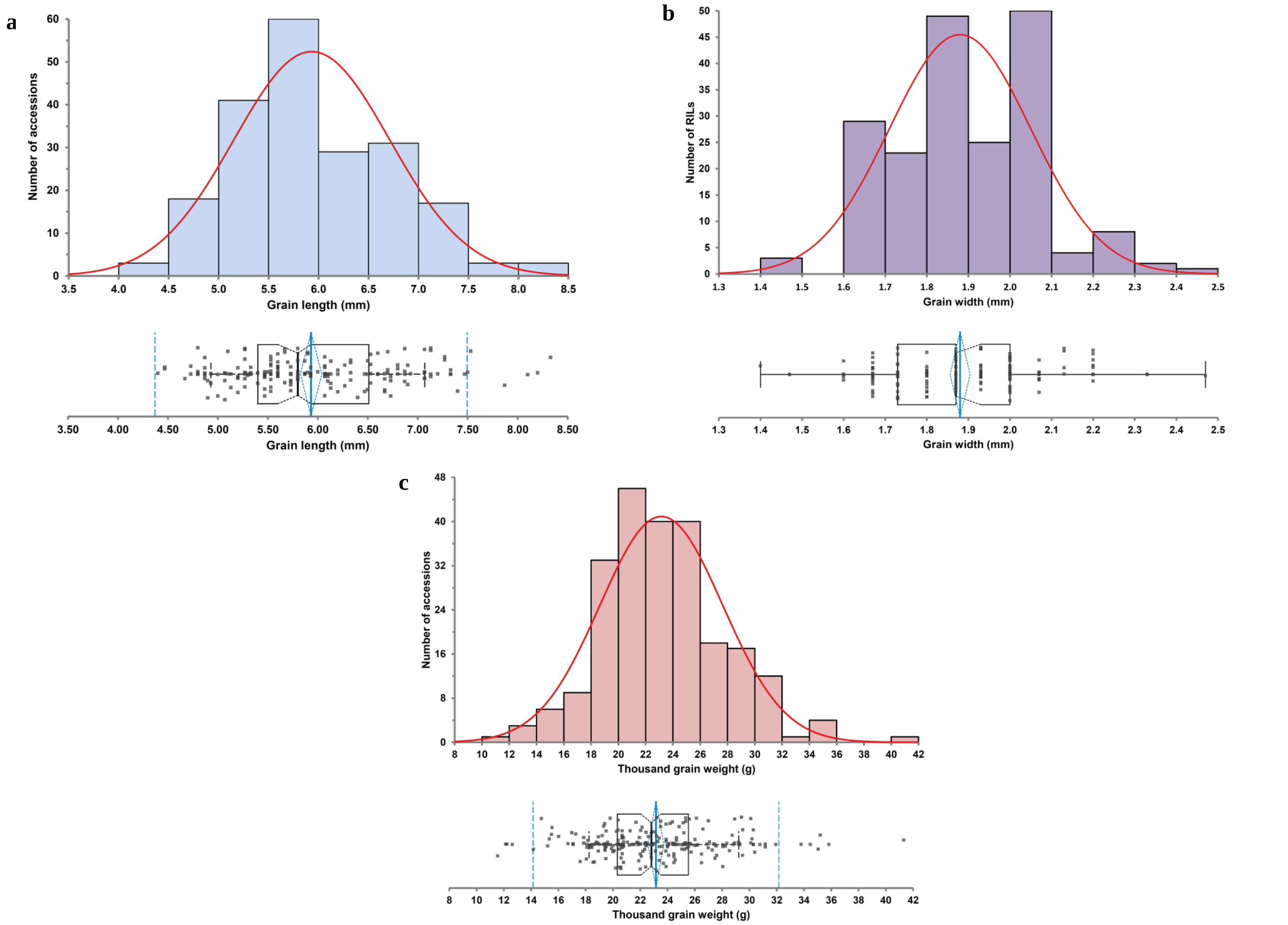


**Supplementary Figure 3. Quantitative genetic inheritance pattern of grain size/weight traits measured in 203 accessions.** Histograms and boxplots depicting distribution of **a)** grain length, **b)** grain width, and **c)** thousand-grain weight distribution in 203 accessions selected from a diversity panel consisting of 271 diverse rice accessions.
