## Supplemental Figure 5 for "Rice Pan-genome Array (RPGA): an efficient genotyping solution for pan-genome-based accelerated crop improvement in rice"

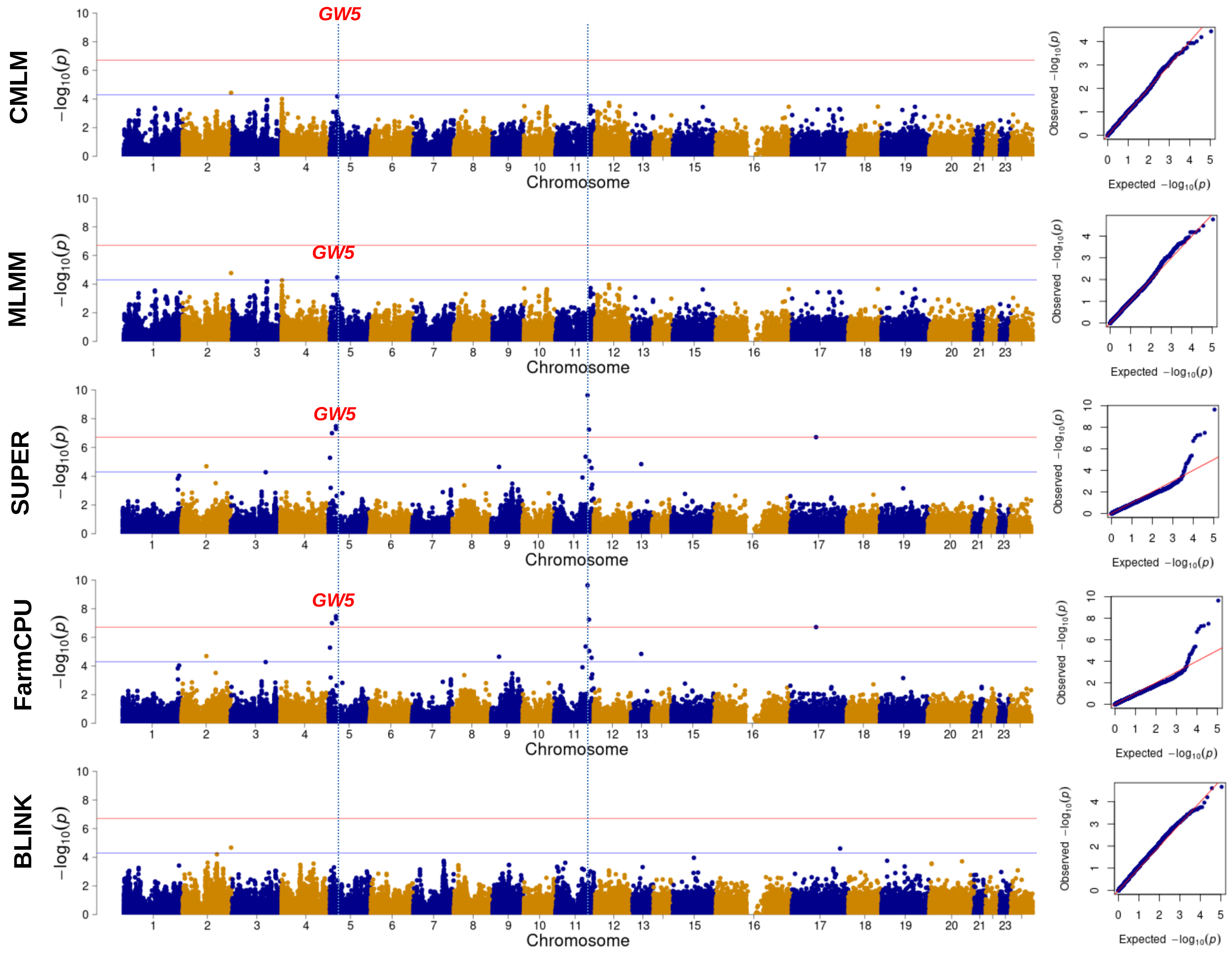


**Supplementary Figure 5. Manhattan plots and QQ plots depicting results of GWAS for grain width trait using five different association models.** Chromosomes 1 to 12 represent twelve Nipponbare chromosomes, whereas, chromosomes 13 to 24 represent twelve sub-population groups specific pseudo-chromosomes from 3K pan-genome. The associated loci that harbor previously known grain width genes are marked on the top. The green horizontal line represents a stringent Benjamini-Hochberg threshold whereas the red horizontal line represents a less stringent suggestive threshold.
