## Supplemental Figure 7 for "Rice Pan-genome Array (RPGA): an efficient genotyping solution for pan-genome-based accelerated crop improvement in rice"

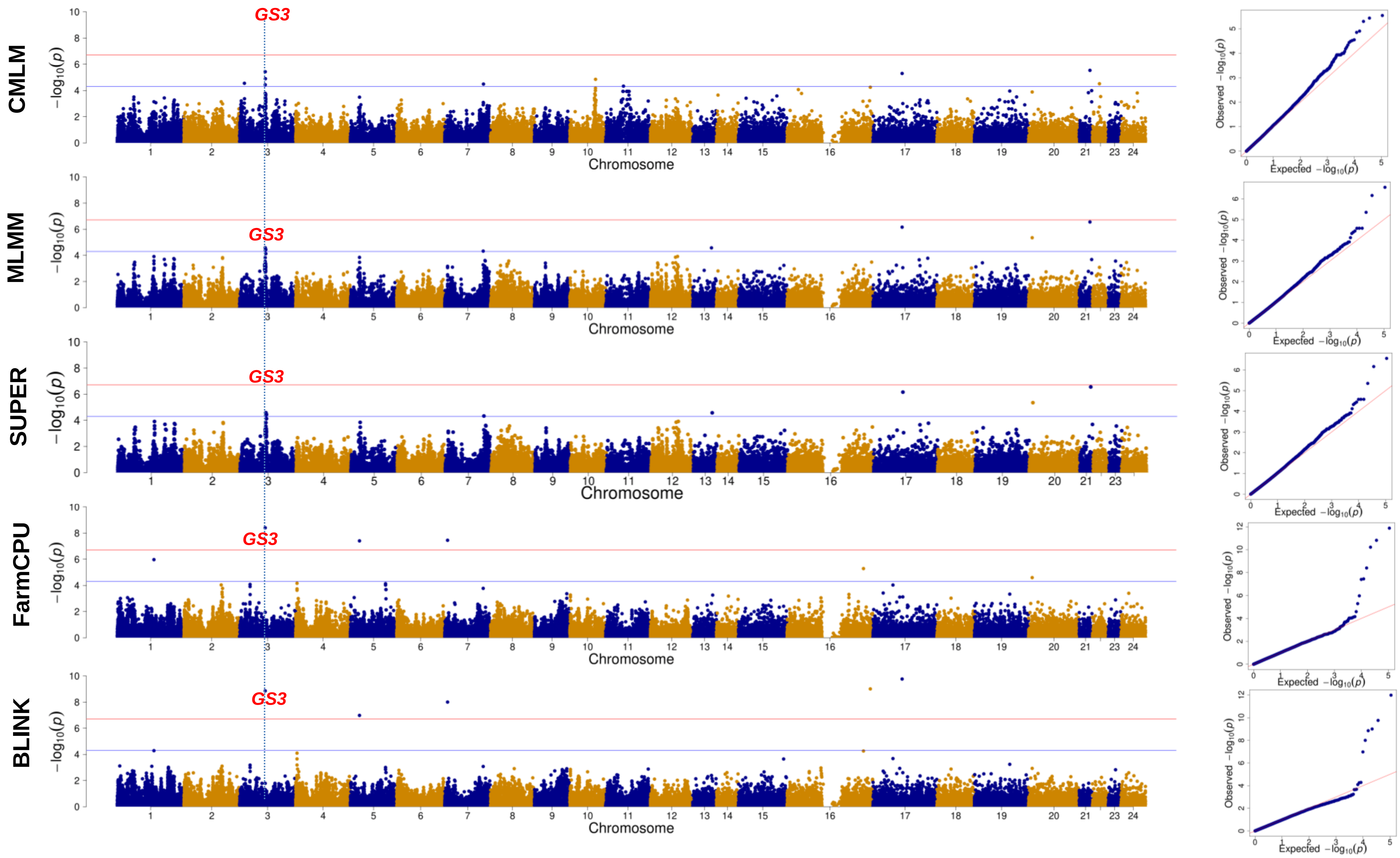


**Supplementary Figure 7. Manhattan plots and QQ plots depicting results of genome-wide association study (GWAS) for length-to-width ratio trait using five different association models**. Chromosomes 1 to 12 represent twelve Nipponbare chromosomes, whereas, chromosomes 13 to 24 represent twelve subpopulation group specific pseudo-chromosomes associated loci which harbor previously known grain weight genes are marked on the top. The blue horizontal line represents a stringent Benjamini-Hochberg threshold whereas the red horizontal line represents a less stringent suggestive threshold.
