## Supplemental Figure 8 for "Rice Pan-genome Array (RPGA): an efficient genotyping solution for pan-genome-based accelerated crop improvement in rice"

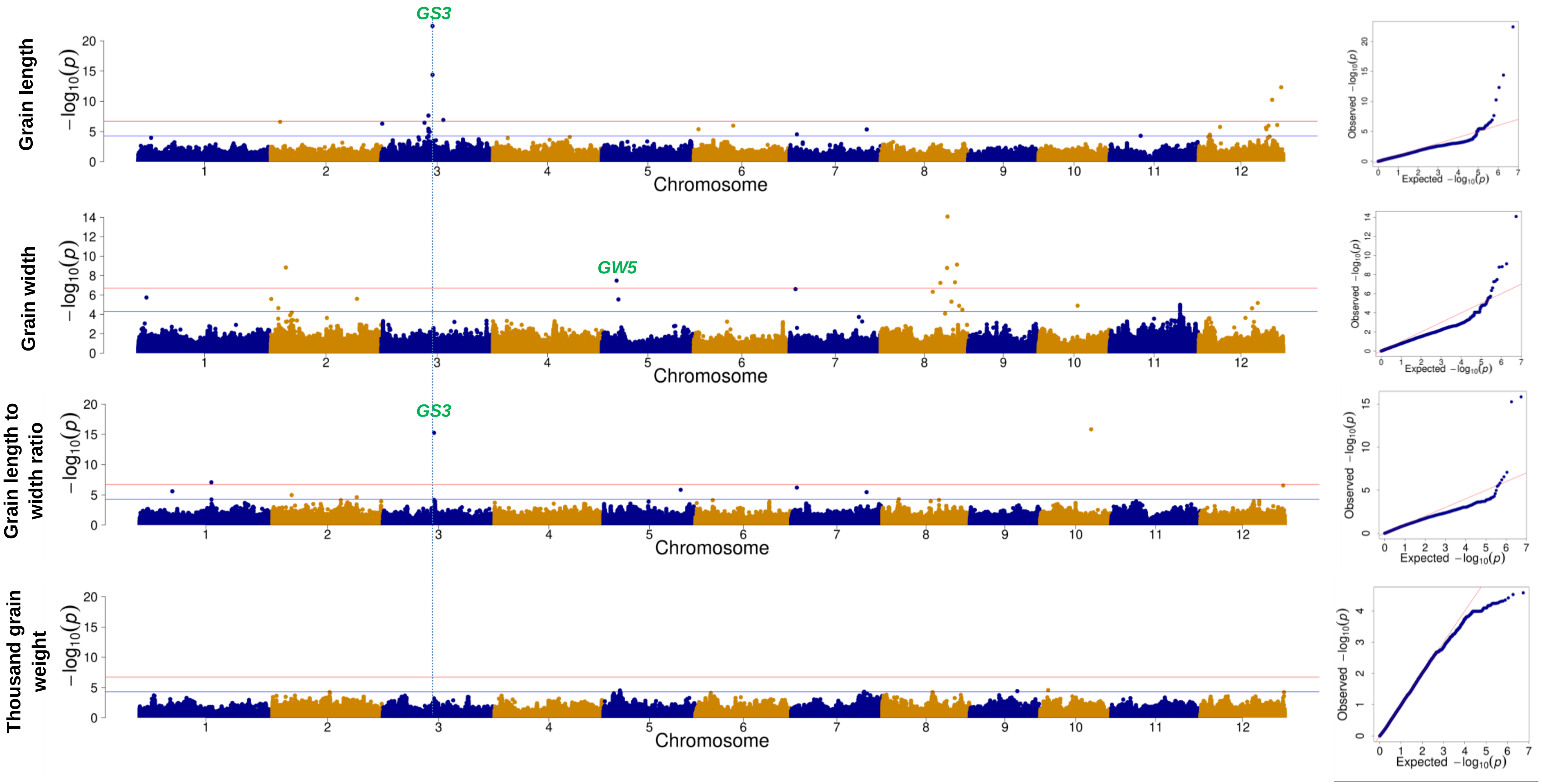


**Supplementary Figure 8. Manhattan plots and QQ plots depicting results of genome-wide association study (GWAS) for grain length, grain width, length-to-width ratio and thousand-grain weight trait performed using imputed genotype data**. Chromosomes 1 to 12 represent twelve Nipponbare chromosomes. Associated loci harboring previously known grain weight genes are marked on the top in green. The blue horizontal line represents a stringent Benjamini-Hochberg threshold whereas the red horizontal line represents a less stringent suggestive threshold.
