## Supplemental Figure 9 for "Rice Pan-genome Array (RPGA): an efficient genotyping solution for pan-genome-based accelerated crop improvement in rice"

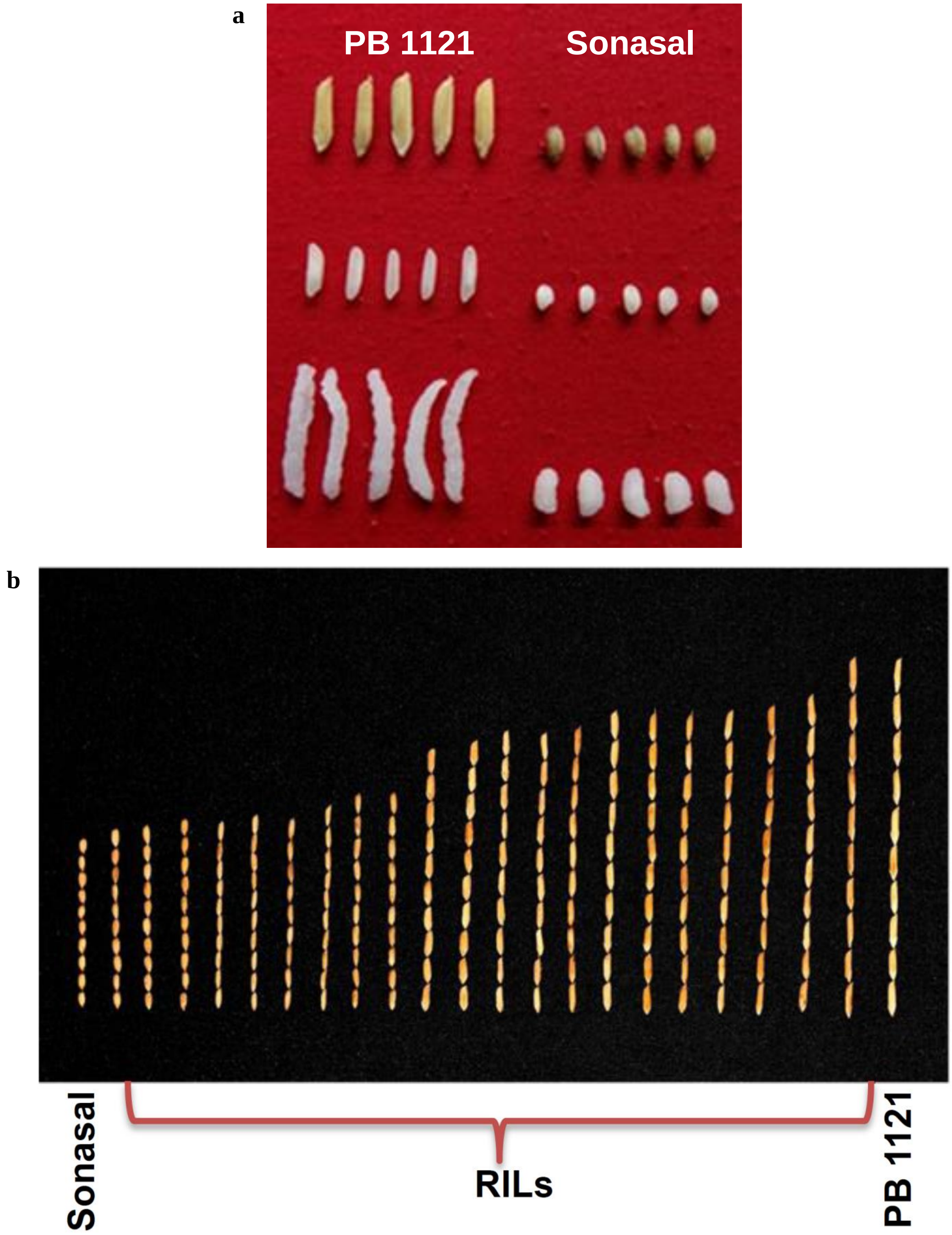


**Supplementary Figure 9. Images depicting grain size variation in a developed mapping population (Sonasal × Pusa Basmati 1121). a)** Rough and milled grains (before and after cooking) of two parental accessions, Sonasal and Pusa Basmati 1121 (PB 1121). **b)** Rough grain variation in a representative set of mapping individuals of a RIL population along with parental accessions, Sonasal and PB 1121.
