## Supplemental Figure 10 for "Rice Pan-genome Array (RPGA): an efficient genotyping solution for pan-genome-based accelerated crop improvement in rice"

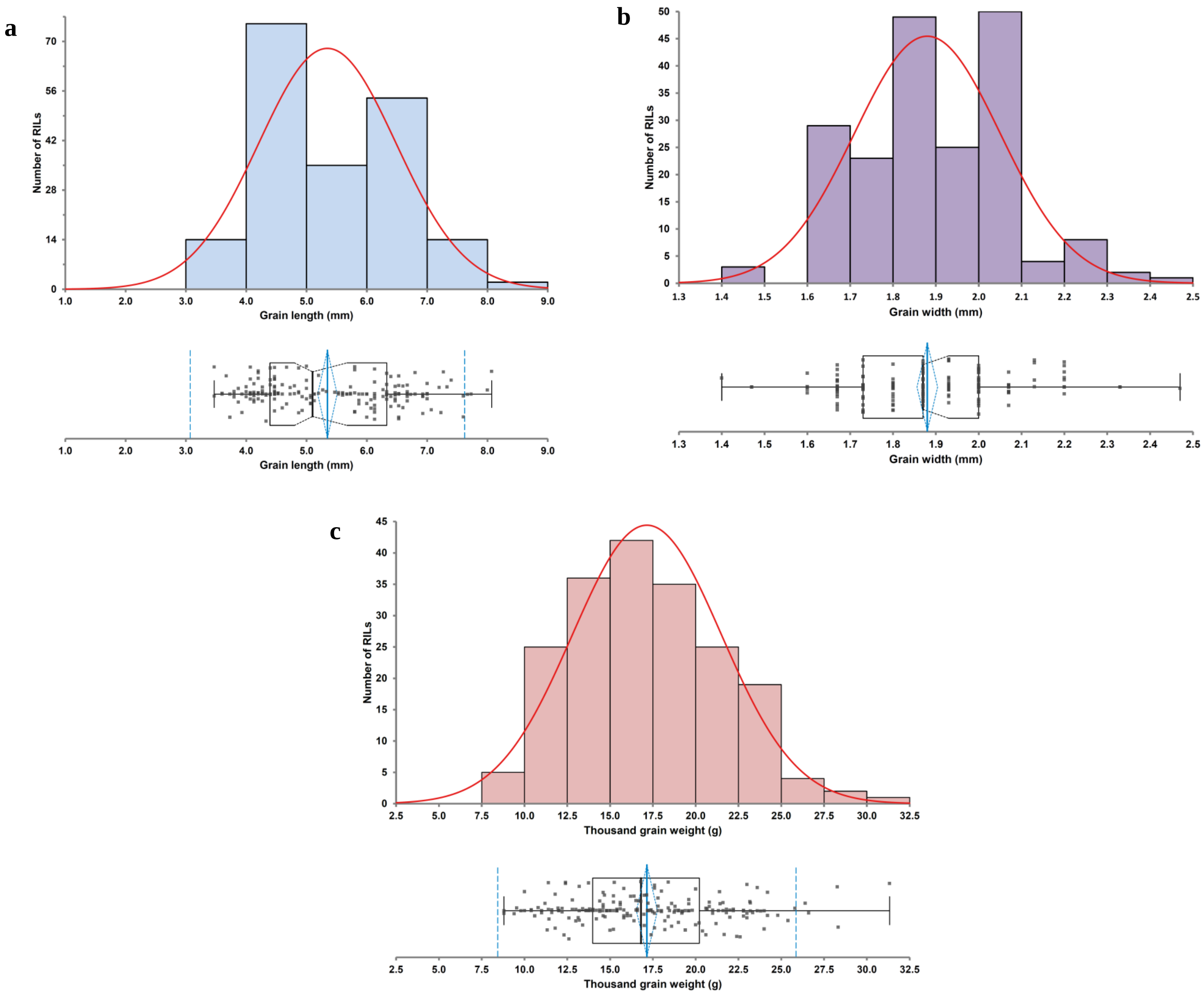


**Supplementary Figure 10. Quantitative genetic inheritance pattern of grain size/weight traits measured in a RIL (Sonsal × Pusa Basmati 1121) mapping population**. Histograms and boxplots depicting distribution of a) grain length, b) grain width, and c) thousand-grain weight in 188 RILs.
