## Supplemental Figure 11 for "Rice Pan-genome Array (RPGA): an efficient genotyping solution for pan-genome-based accelerated crop improvement in rice"

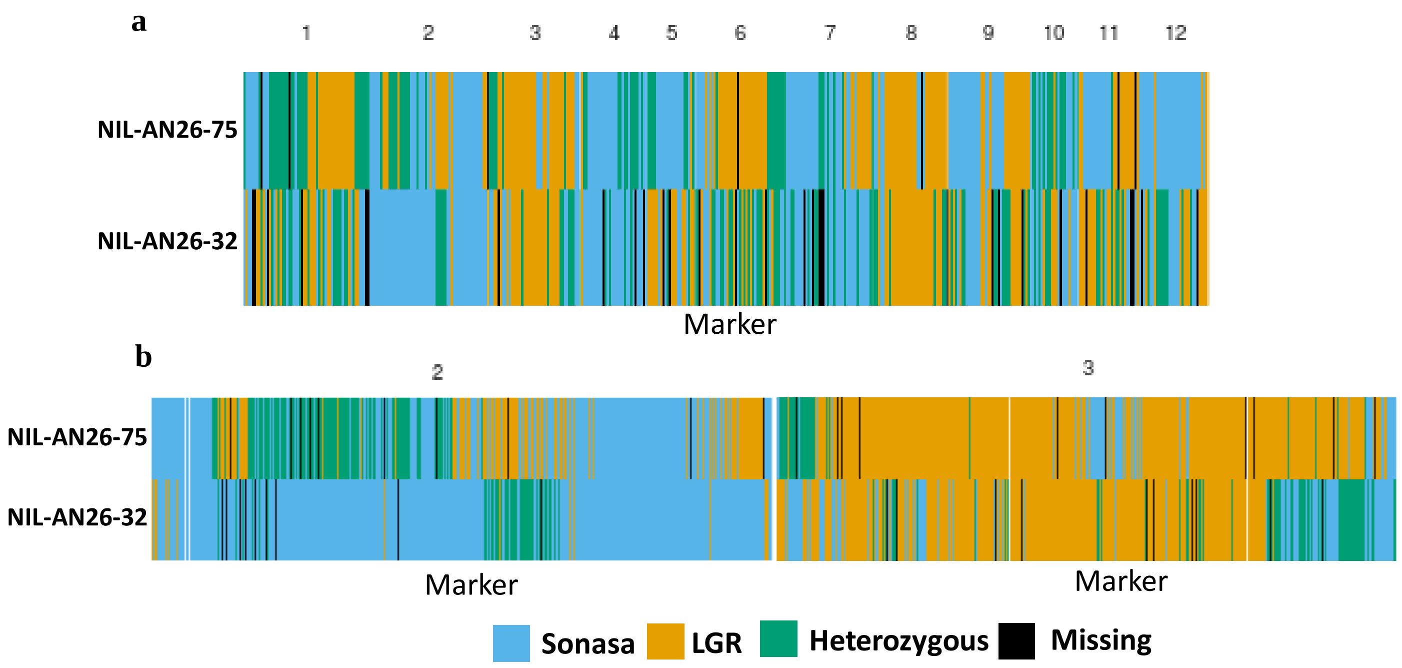


**Supplementary Figure 11. Background analysis of two Sonasal × LGR $BC_4F_1$ lines. a)** Graphical of Genotype of all 12 rice chromosome, and **b)** Graphical of Genotype of chromosome 2 and chromosome 3.
