## Supplementary figures and images for "Rice Pan-genome Array (RPGA): an efficient genotyping solution for pan-genome-based accelerated crop improvement in rice"

### Supplemental Figure 12

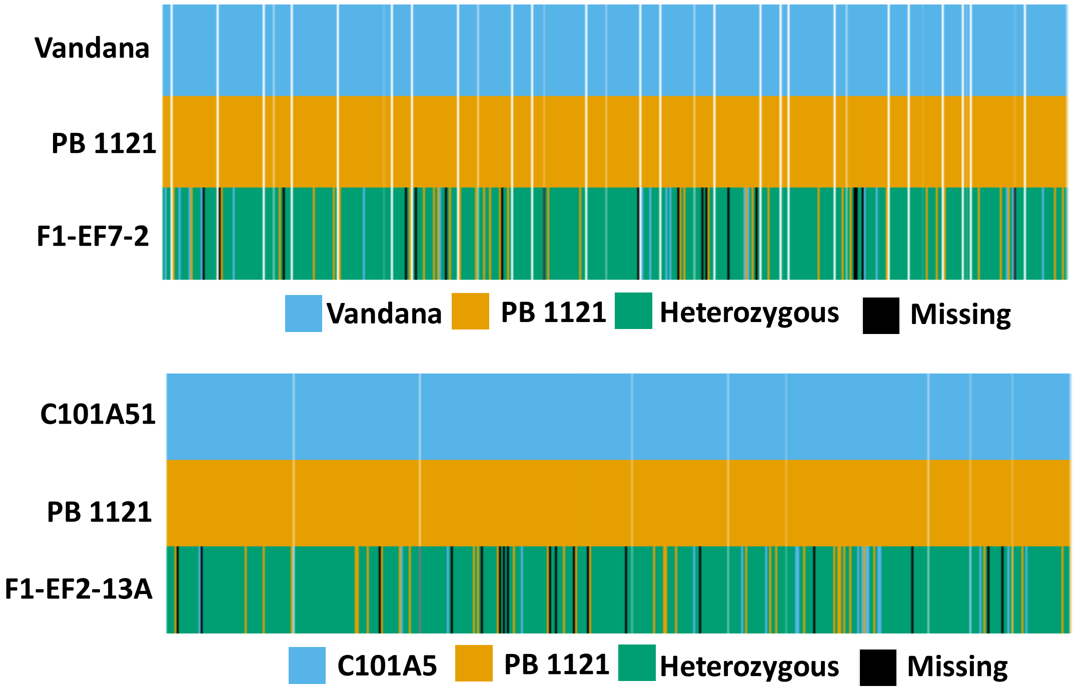


**Supplementary Figure 12. Hybridity test of $F_1$ hybrids performed using RPGA-based genotyping.**
