## Supplemental Figure 13 for "Rice Pan-genome Array (RPGA): an efficient genotyping solution for pan-genome-based accelerated crop improvement in rice"

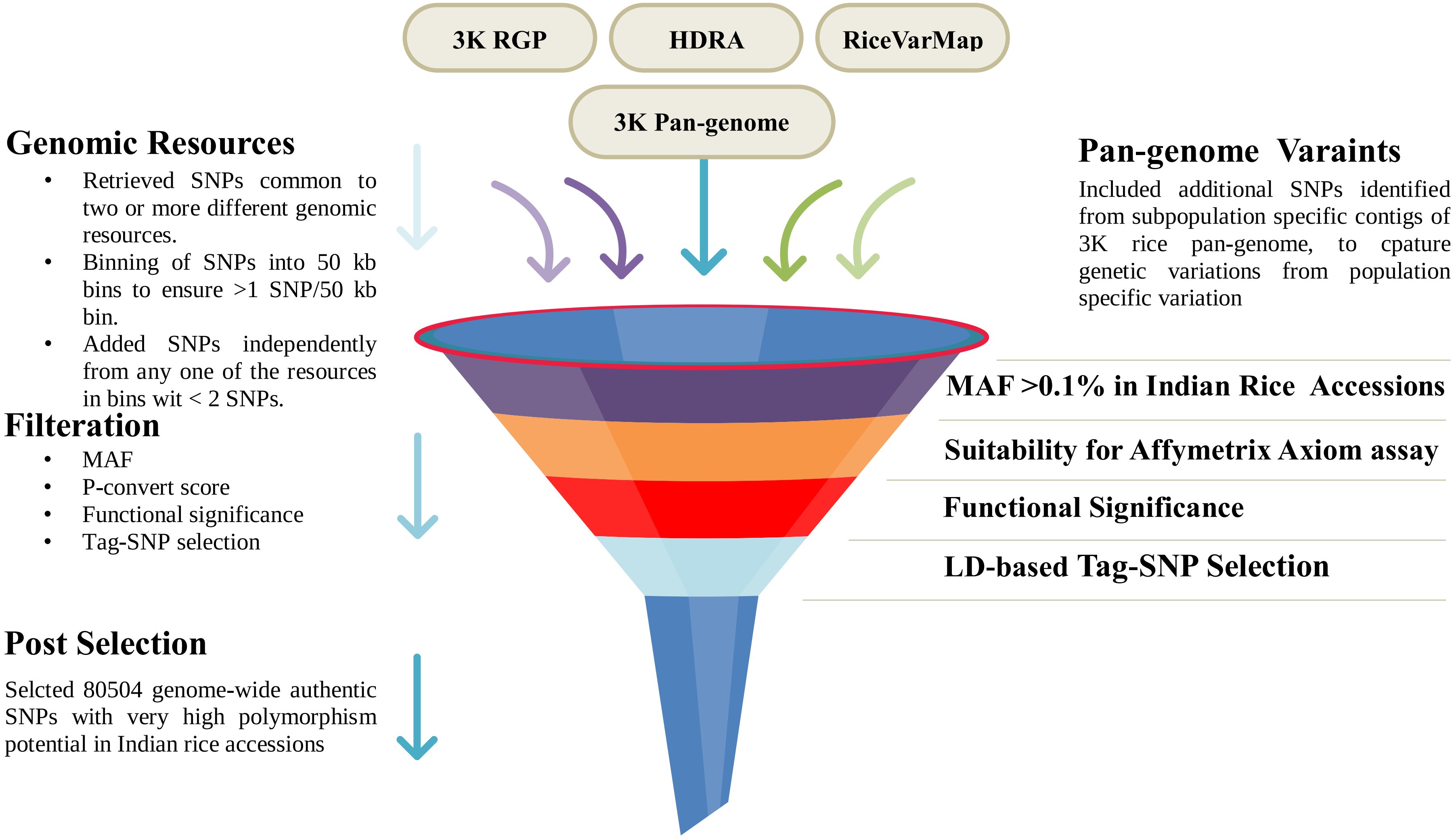


**Supplementary Figure 13. Workflow describing steps involved in designing of Rice Pan-genome Genome Genotyping Array (RPGA)**. 3K RGP: 3K Rice Genomes Project, HDRA: High-Density Rice Array, MAF: Minor Allele Frequency
