## Supplemental Table 1 for "Rice Pan-genome Array (RPGA): an efficient genotyping solution for pan-genome-based accelerated crop improvement in rice"

**Supplementary Table 1.** Classification of SNPs into six distinct classes produced from RIL genotyping data by Affymetrix Axiom best practice genotyping workflow

| **Conversion Type** | **Count** | **Percentage** |
|---|---|---|
| Poly High Resolution (PHR) | 46523 | 57.78 |
| Other | 11468 | 14.25 |
| Mono High Resolution (MHR) | 9013 | 11.20 |
| No Minor Homozygous (NMH) | 6841 | 8.50 |
| Off Target Variants (OTV) | 5746 | 7.14 |
| Call Rate Below Threshold (CRBT) | 913 | 1.13 |
