## Supplemental Table 2 for "Rice Pan-genome Array (RPGA): an efficient genotyping solution for pan-genome-based accelerated crop improvement in rice"

**Supplementary Table 2.** Distribution of SNPs in six different classes from genotyping data of a diversity panel

| **Conversion Type** | **Count** | **Percentage** |
|---|---|---|
| Poly High Resolution (PHR) | 48133 | 59.79 |
| Off-target variants (OTV) | 12618 | 15.67 |
| Other | 12160 | 15.11 |
| No Minor Homozygous (NMH) | 2646 | 3.29 |
| Call Rate Below Threshold (CRBT) | 2645 | 3.29 |
| Mono High Resolution (MHR) | 2302 | 2.86 |
