## Supplemental Table 3 for "Rice Pan-genome Array (RPGA): an efficient genotyping solution for pan-genome-based accelerated crop improvement in rice"

**Supplementary Table 3.** Characteristics of an ultra-density-genetic map developed using SNPs annotated on the Nipponbare genome

| **Chromosomes** | **Number of Markers** | **Length (cM)** | **Average Spacing (cM)** | **Maximum Spacing (cM)** |
|---|---|---|---|---|
| 1 | 1596 | 179.4 | 0.1 | 8.3 |
| 2 | 848 | 129.5 | 0.2 | 17.3 |
| 3 | 850 | 148.7 | 0.2 | 9.1 |
| 4 | 723 | 112.4 | 0.2 | 7.7 |
| 5 | 1073 | 111.7 | 0.1 | 12.3 |
| 6 | 689 | 140.7 | 0.2 | 41.9 |
| 7 | 902 | 82.6 | 0.1 | 12.1 |
| 8 | 946 | 84.7 | 0.1 | 10.5 |
| 9 | 252 | 93.8 | 0.4 | 37.1 |
| 10 | 559 | 93.3 | 0.2 | 8.2 |
| 11 | 678 | 122.2 | 0.2 | 42.2 |
| 12 | 747 | 117.5 | 0.2 | 37.3 |
| **Total** | **9863** | **1416.4** | **0.1** | **42.2** |
