## Supplemental Table 4 for "Rice Pan-genome Array (RPGA): an efficient genotyping solution for pan-genome-based accelerated crop improvement in rice"

**Supplementary Table 4.** Characteristics of an ultra-density-genetic map developed using combining SNPs annotated on the Nipponbare genome as well as 12 population-specific pseudo-chromosomes

| **Chromosomes** | **Number of Markers** | **Length (cM)** | **Average Spacing (cM)** | **Maximum Spacing (cM)** |
|---|---|---|---|---|
| 1 | 2035 | 311.5 | 0.2 | 13.5 |
| 2 | 1146 | 200.2 | 0.2 | 17.5 |
| 3 | 991 | 277.8 | 0.3 | 11.4 |
| 4 | 1037 | 214.6 | 0.2 | 8.3 |
| 5 | 1403 | 259.4 | 0.2 | 14.8 |
| 6 | 916 | 159.6 | 0.2 | 21.3 |
| 7 | 1315 | 145 | 0.1 | 16.6 |
| 8 | 1241 | 197.3 | 0.2 | 9.5 |
| 9 | 329 | 109.2 | 0.3 | 22.2 |
| 10 | 1156 | 153.9 | 0.1 | 6.8 |
| 11 | 980 | 137.6 | 0.1 | 7.4 |
| 12 | 1244 | 146.2 | 0.1 | 12.6 |
| **Total** | **13793** | **2312.2** | **0.2** | **22.2** |
