## Supplemental Table 5 for "Rice Pan-genome Array (RPGA): an efficient genotyping solution for pan-genome-based accelerated crop improvement in rice"

**Supplementary Table 5.** Details of GWAS for grain length, grain width and thousand-grain weight trait performed using CMLM

| SNP-ID | Chr/Pseudo-chr | Physical position (bp) | Grain length | | Grain width | | Length-to-width ratio | | Thousand-grain weight | | Structural annotation* | Candidate genes |
|---|---|---|---|---|---|---|---|---|---|---|---|---|
| | | | P-value | FDR adjusted P-values | P-value | FDR adjusted P-values | P-value | FDR adjusted P-values | P-value | FDR adjusted P-values | | |
| JG10~6950498 | JG10 | 6950498 | 1.07E-11 | 5.87E-07 | NA | NA | NA | NA | NA | NA | NA | *WD repeat-containing protein 12* (*LOC_Os07g4093*) |
| JG10~4245875 | JG10 | 4245875 | 3.38E-08 | 0.001 | NA | NA | NA | NA | NA | NA | NA | Unknown function (*OsR498G0612839600*) |
| 7~25105073 | Chr7 | 25105073 | 2.15E-07 | 0.004 | NA | NA | NA | NA | NA | NA | Upstream | *Leaf and flower related protein* (*LOC_Os07g41900*) |
| 3~16733441 | Chr3 | 16733441 | 2.86E-07 | 0.004 | NA | NA | NA | NA | NA | NA | Intergenic | *GS3* (*Os03g0407400*)# |
| 12~25514661 | Chr12 | 25514661 | 8.68E-07 | 0.008 | NA | NA | NA | NA | NA | NA | Intergenic | *IQ calmodulin-binding motif family protein* (LOC_Os12g41160) |
| 1~34538998 | Chr1 | 34538998 | 8.74E-07 | 0.008 | NA | NA | NA | NA | NA | NA | Intergenic | *qGRL1.1/OsGAMYB* (*LOC_Os01g59660*) |
| 11~6484221 | Chr11 | 6484221 | 3.51E-06 | 0.027 | NA | NA | NA | NA | NA | NA | Upstream | NA |
| 10~20357189 | Chr10 | 20357189 | 8.13E-06 | 0.055 | NA | NA | NA | NA | NA | NA | | NA |
| JG10~6243872 | JG10 | 6243872 | 2.46E-05 | 0.149 | NA | NA | NA | NA | 5.69E-08 | 0.0020 | NA | NA |
| 11~24534050 | Chr11 | 24534050 | NA | NA | 2.35E-10 | 1.29E-05 | NA | NA | NA | NA | 3′ UTR | *Receptor-like protein kinase* (*LOC_Os11g4097*) |
| 5~5371686 | Chr5 | 5371686 | NA | NA | 3.36E-08 | 0.00078 | NA | NA | NA | NA | Downstrem | *GW5* (*LOC_Os05g0952*) |
| 11~25612932 | Chr11 | 25612932 | NA | NA | 5.73E-08 | 0.00078 | NA | NA | NA | NA | Intergenic | NA |
| 5~2422324 | Chr5 | 2422324 | NA | NA | 1.01E-07 | 0.001 | NA | NA | NA | NA | 3′ UTR | NA |
| IG2~18885542 | IG2 | 18885542 | NA | NA | 1.93E-07 | 0.001 | NA | NA | NA | NA | NA | *LOC_Os10g31770* |
| 11~23154487 | Chr11 | 23154487 | NA | NA | 4.38E-06 | 0.034 | NA | NA | NA | NA | Synonymous | NA |
| 5~991259 | Chr5 | 991259 | NA | NA | 5.25E-06 | 0.035 | NA | NA | NA | NA | 5′ UTR | NA |

| SNP-ID | Chr/Pseudo-chr+ | Physical position (bp) | Grain length | | Grain width | | Length-to-width ratio | | Thousand-grain weight | | Structural annotation* | Candidate genes |
|---|---|---|---|---|---|---|---|---|---|---|---|---|
| | | | P-value | FDR adjusted P-values | P-value | FDR adjusted P-values | P-value | FDR adjusted P-values | P-value | FDR adjusted P-values | | |
| 11~25783593 | Chr11 | 25783593 | NA | NA | 8.95E-06 | 0.054 | NA | NA | NA | NA | Intergenic | NA |
| Adm~7425964 | Adm | 7425964 | NA | NA | 1.44E-05 | 0.078 | NA | NA | NA | NA | NA | NA |
| 9~5770786 | Chr9 | 5770786 | NA | NA | 2.25E-05 | 0.102 | NA | NA | NA | NA | Intergenic | NA |
| 11~27490883 | Chr11 | 27490883 | NA | NA | 2.65E-05 | 0.111 | NA | NA | NA | NA | Downstream | NA |
| 3~25711563 | Chr3 | 25711563 | NA | NA | 5.30E-05 | 0.207 | NA | NA | NA | NA | Intergenic | NA |
| 5~5390763 | Chr5 | 5390763 | NA | NA | NA | NA | 2.30E-05 | 0.7946 | 7.60E-08 | 0.00207 | Downstream | *GW5* (*LOC_Os05g09520*) |
| 10~21588761 | Chr10 | 21588761 | NA | NA | NA | NA | NA | NA | 4.14E-05 | 0.454 | Intergenic | NA |
| 11~2611306 | Chr11 | 2611306 | NA | NA | NA | NA | NA | NA | 4.15E-05 | 0.454 | Upstream | NA |
| 8~17187746 | Chr8 | 17187746 | NA | NA | NA | NA | NA | NA | 5.82E-05 | 0.501 | Upstream | NA |
| IG5~14354843 | IG5 | 14354843 | 2.90E-05 | NA | NA | NA | 2.90E-05 | 0.7946 | NA | NA | NA | |
| IG5~14354791 | IG5 | 14354791 | 5.26E-05 | NA | NA | NA | 5.26E-05 | 0.9610 | NA | NA | NA | |
| 5~4222429 | Chr5 | 4222429 | NA | NA | NA | NA | NA | NA | 6.42E-05 | 0.501 | Intergenic | NA |

Chr: Chromosome; Pseudo-chr: Pseudo-chromosome; 5′ UTR/3′ UTR: five prime untranslated region/ three prime untranslated region defined as per MSU7.0 annotation; Upstream: SNPs located >2 kb upstream from transcription start site of a gene; Downstream: SNPs located >2 kb downstream from transcription stop site of a gene; IG: Indica Group; JG: Japonica Group; Adm: Admixture *Structural annotation performed using MSU 7.0 ; #Rice Annotation Project locus ID; The loci detected from pseudo-chromosomes are highlighted in red.
