## Supplemental Table 6 for "Rice Pan-genome Array (RPGA): an efficient genotyping solution for pan-genome-based accelerated crop improvement in rice"

**Supplementary Table 6.** Details of GWAS for grain length, grain width and thousand-grain weight trait performed using MLMM

| SNP-ID | Chr/Pseudo-chr+ | Physical position (bp) | Grain length | | Grain width | | Length-to-width ratio | | Thousand-grain weight | | Structural annotation* | Candidate genes |
|---|---|---|---|---|---|---|---|---|---|---|---|---|
| | | | P-value | FDR adjusted P-values | P-value | FDR adjusted P-values | P-value | FDR adjusted P-values | P-value | FDR adjusted P-values | | |
| 3~16733441 | Chr3 | 16733441 | 1.66E-08 | 0.00091 | NA | NA | NA | NA | NA | NA | Intergenic | *GS3 (Os03g0407400)* |
| 3~3775429 | Chr3 | 3775429 | 2.29E-06 | 0.06276 | NA | NA | NA | NA | NA | NA | Upstream | *Positve Regulator of Grain Length 1 (LOC_Os03g07510)* |
| 3~34617712 | Chr3 | 34617712 | 1.69E-05 | 0.18456 | NA | NA | NA | NA | NA | NA | Upstream | NA |
| 4~11470427 | Chr4 | 11470427 | 1.76E-05 | 0.18456 | NA | NA | NA | NA | NA | NA | Upstream | NA |
| 4~1881177 | Chr4 | 1881177 | 2.28E-05 | 0.18456 | NA | NA | NA | NA | NA | NA | Intergenic | NA |
| 3~34643467 | Chr3 | 34643467 | 2.69E-05 | 0.18456 | NA | NA | NA | NA | NA | NA | Downstream | NA |
| 7~25053326 | Chr7 | 25053326 | 3.03E-05 | 0.18456 | NA | NA | NA | NA | NA | NA | Downstream | NA |
| 8~16338601 | Chr8 | 16338601 | 3.44E-05 | 0.18456 | NA | NA | NA | NA | NA | NA | Intergenic | NA |
| JG8~4694243 | JG8 | 4694243 | 3.79E-05 | 0.18456 | NA | NA | NA | NA | NA | NA | NA | NA |
| AUSG6~24889549 | AUSG6 | 24889549 | 4.04E-05 | 0.18456 | NA | NA | NA | NA | NA | NA | NA | NA |
| JG10~6950498 | JG10 | 6950498 | 5.13E-05 | 0.18456 | NA | NA | NA | NA | NA | NA | NA | NA |
| JG9~3630875 | JG9 | 3630875 | 5.94E-05 | 0.18456 | NA | NA | NA | NA | NA | NA | NA | NA |
| 2~35350740 | Chr2 | 35350740 | NA | NA | 1.69E-05 | 2~35350740 | NA | NA | NA | NA | Intergenic | NA |
| 5~5371686 | Chr5 | 5371686 | NA | NA | 3.31E-05 | 5~5371686 | NA | NA | NA | NA | Downstream | *GW5 (LOC_Os05g09520)* |
| 4~363030 | Chr4 | 363030 | NA | NA | 5.40E-05 | 4~363030 | NA | NA | NA | NA | 3′ UTR | NA |
| 3~25782328 | Chr3 | 25782328 | NA | NA | 6.60E-05 | 3~25782328 | NA | NA | NA | NA | Intergenic | NA |
| 5~5390763 | 5 | 5390763 | NA | NA | NA | NA | 9.882E-06 | 0.348 | NA | NA | Downstream | *GW5* (*LOC_Os05g09520*) |
| IG5~14354843 | IG5 | 14354843 | NA | NA | NA | NA | 1.27E-05 | 0.3482 | 1.27E-05 | 0.348 | Downstream | *GW 5 (LOC_Os05g09520)* |
| IG5~14354791 | IG5 | 14354791 | NA | NA | NA | NA | 2.645E-05 | 0.48256 | 2.65E-05 | 0.483 | NA | NA |

Chr: Chromosome; Pseudo-chr: Pseudo-chromosome; 5′ UTR/3′ UTR: five prime untranslated region/ three prime untranslated region defined as per MSU7.0 annotation; Upstream: SNPs located >2 kb upstream from transcription start site of a gene; Downstream: SNPs located >2 kb downstream from transcription stop site of a gene; IG: Indica Group; JG: Japonica Group; Adm: Admixture *Structural annotation performed using MSU 7.0; #Rice Annotation Project locus ID; The loci detected from pseudo-chromosomes are highlighted in red.
