## Supplemental Table 7 for "Rice Pan-genome Array (RPGA): an efficient genotyping solution for pan-genome-based accelerated crop improvement in rice"

**Supplementary Table 7.** Details of GWAS for grain length, grain width and thousand-grain weight trait performed using SUPER

| SNP-ID | Chr/Pseudo-chr+ | Physical position (bp) | Grain length | | Grain width | | Length-to-width ratio | | Thousand-grain weight | | Structural annotation* | Candidate genes |
|---|---|---|---|---|---|---|---|---|---|---|---|---|
| | | | P-value | FDR adjusted P-values | P-value | FDR adjusted P-values | P-value | FDR adjusted P-values | P-value | FDR adjusted P-values | | |
| JG10~6950498 | JG10 | 6950498 | 1.07E-11 | 5.87E-07 | NA | NA | NA | NA | NA | NA | NA | *WD repeat-containing protein 12 (LOC_Os07g40930)* |
| JG10~4245875 | JG10 | 4245875 | 3.38E-08 | 0.000925 | NA | NA | NA | NA | NA | NA | NA | *Unknown function (OsR498G0612839600.01)* |
| 7~25105073 | Chr7 | 2.5E+07 | 2.15E-07 | 0.003919 | NA | NA | NA | NA | NA | NA | Upstream | *Leaf and flower related protein (LOC_Os07g41900)* |
| 3~16733441 | Chr3 | 1.7E+07 | 2.86E-07 | 0.003919 | NA | NA | NA | NA | NA | NA | Intergenic | *GS3 (Os03g0407400)* |
| 12~25514661 | Chr12 | 2.6E+07 | 8.68E-07 | 0.007971 | NA | NA | NA | NA | NA | NA | Intergenic | *IQ calmodulin-binding motif family protein ( LOC_Os12g41160)* |
| 1~34538998 | Chr1 | 3.5E+07 | 8.74E-07 | 0.007971 | NA | NA | NA | NA | NA | NA | Intergenic | *qGRL1.1/OsGAMYB (LOC_Os01g59660)* |
| 11~6484221 | Chr11 | 6484221 | 3.51E-06 | 0.027446 | NA | NA | NA | NA | NA | NA | Upstream | NA |
| 10~20357189 | Chr10 | 2E+07 | 8.13E-06 | 0.055605 | NA | NA | NA | NA | NA | NA | Intergenic | NA |
| JG10~6243872 | JG10 | 6243872 | 2.46E-05 | 0.149332 | NA | NA | 5.69E-08 | 0.0020 | 5.69E-08 | 0.0020783 | NA | NA |
| 11~24534050 | Chr11 | 2.5E+07 | NA | NA | 2.35E-10 | 1.29E-05 | NA | NA | NA | NA | 3′ UTR | *Receptor-like protein kinase precursor (LOC_Os11g40970)* |
| 5~5371686 | Chr5 | 5371686 | NA | NA | 3.36E-08 | 0.0007 | NA | NA | NA | NA | Downstream | *GW5 (LOC_Os05g09520)* |
| 5~5390763 | Chr5 | 5390763 | NA | NA | NA | NA | 0.00000007 | 0.348 | NA | NA | Downstream | *GW5* (*LOC_Os05g09520*) |
| IG2~18885542 | IG2 | 1.9E+07 | NA | NA | 1.93E-07 | 0.001756 | NA | NA | NA | NA | NA | *LOC_Os10g31770* |

| SNP-ID | Chr/Pseudo-chr+ | Physical position (bp) | Grain length | | Grain width | | Length-to-width ratio | | Thousand-grain weight | | Structural annotation* | Candidate genes |
|---|---|---|---|---|---|---|---|---|---|---|---|---|
| | | | P-value | FDR adjusted P-values | P-value | FDR adjusted P-values | P-value | FDR adjusted P-values | P-value | FDR adjusted P-values | | |
| 11~23154487 | Chr11 | 2.30E+07 | NA | NA | 4.38E-06 | 0.0342 | NA | NA | NA | NA | Synonymous | NA |
| 5~991259 | Chr5 | 991259 | NA | NA | 5.25E-06 | 0.0359 | NA | NA | NA | NA | 5′ UTR | NA |
| 11~25783593 | Chr11 | 2.60E+07 | NA | NA | 8.95E-06 | 0.0544 | NA | NA | NA | NA | Intergenic | NA |
| Adm~7425964 | Adm | 7425964 | NA | NA | 1.44E-05 | 0.0789 | NA | NA | NA | NA | NA | NA |
| 2~18141348 | Chr2 | 1.80E+07 | NA | NA | 2.03E-05 | 0.1011 | NA | NA | NA | NA | Intergenic | NA |
| 9~5770786 | Chr9 | 5770786 | NA | NA | 2.25E-05 | 0.1027 | NA | NA | NA | NA | Intergenic | NA |
| 11~27490883 | Chr11 | 2.70E+07 | NA | NA | 2.65E-05 | 0.1116 | NA | NA | NA | NA | Downstream | NA |
| 3~25711563 | Chr3 | 2.60E+07 | NA | NA | 5.30E-05 | 0.2073 | NA | NA | NA | NA | Intergenic | NA |
| 5~5390763 | Chr5 | 5390763 | NA | NA | NA | NA | 7E-08 | 0.348 | 7.60E-08 | 0.002 | Downstream | *GW5 (LOC_Os05g09520)* |
| 4~4486000 | Chr4 | 4486000 | NA | NA | NA | NA | 7.98E-06 | 0.145 | 7.99E-06 | 0.145 | Missense | *LRR family protein encoding gene* (*LOC_Os04g08390*) |
| 10~21588761 | Chr10 | 2.20E+07 | NA | NA | NA | NA | 4.14E-05 | 0.454 | 4.14E-05 | 0.454 | Intergenic | NA |
| 11~2611306 | Chr11 | 2611306 | NA | NA | NA | NA | 0.00004 | 0.454 | 4.15E-05 | 0.454 | Upstream | NA |
| 8~17187746 | Chr8 | 1.70E+07 | NA | NA | NA | NA | 5.82E-05 | 0.501 | 5.82E-05 | 0.501 | Upstream | NA |
| 5~4222429 | Chr5 | 4222429 | NA | NA | NA | NA | 6.41E-05 | 0.501 | 6.42E-05 | 0.501 | Intergenic | NA |

Chr: Chromosome; Pseudo-chr: Pseudo-chromosome; 5′ UTR/3′ UTR: five prime untranslated region/ three prime untranslated region defined as per MSU7.0 annotation; Upstream: SNPs located >2 kb upstream from transcription start site of a gene; Downstream: SNPs located >2 kb downstream from transcription stop site of a gene; IG: Indica Group; JG: Japonica Group; Adm: Admixture *Structural annotation performed using MSU 7.0; #Rice Annotation Project locus ID; The loci detected from pseudo-chromosomes are highlighted in red.
