## Supplemental Table 9 for "Rice Pan-genome Array (RPGA): an efficient genotyping solution for pan-genome-based accelerated crop improvement in rice"

**Supplementary Table 9.** Details of GWAS for grain length, grain width and thousand-grain weight trait performed using BLINK

| SNP-ID | Chr/Pseudo-chr+ | Physical position (bp) | Grain length | | Grain width | | Length-to-width ratio | | Thousand-grain weight | | Structural annotation* | Candidate genes |
|---|---|---|---|---|---|---|---|---|---|---|---|---|
| | | | P-value | FDR adjusted P-values | P-value | FDR adjusted P-values | P-value | FDR adjusted P-values | P-value | FDR adjusted P-values | | |
| JG10~6950498 | JG10 | 6950498 | 3.25E-10 | 1.78E-05 | NA | NA | NA | NA | NA | NA | NA | *WD repeat-containing protein 12 (LOC_Os07g40930)* |
| 3~3775429 | Chr 3 | 3775429 | 5.46E-08 | 1.49E-03 | NA | NA | NA | NA | NA | NA | Upstream | *PGL1 (LOC_Os03g07510)* |
| 4~1881177 | Chr 4 | 1881177 | 5.73E-06 | 0.104 | NA | NA | NA | NA | NA | NA | Intergenic | NA |
| JG10~4325684 | JG10 | 4325684 | 4.47E-05 | 0.499 | NA | NA | NA | NA | NA | NA | NA | NA |
| AUSG6~24889549 | AUSG6 | 24889549 | 4.57E-05 | 0.499 | NA | NA | NA | NA | NA | NA | NA | NA |
| JG10~2239066 | JG10 | 2239066 | 5.86E-05 | 0.534 | NA | NA | NA | NA | NA | NA | NA | NA |
| 6~2735663 | Chr 6 | 2735663 | 7.72E-05 | 0.603 | NA | NA | NA | NA | NA | NA | Downstream | NA |
| JG10~4882245 | JG10 | 4882245 | 8.86E-05 | 0.606 | NA | NA | NA | NA | NA | NA | NA | NA |
| 2~35350740 | Chr2 | 35350740 | NA | NA | 2.11E-05 | 0.661 | NA | NA | NA | NA | Intergenic | NA |
| IG2~35461875 | Chr17 | 35461875 | NA | NA | 2.44E-05 | 0.661 | NA | NA | NA | NA | NA | NA |
| 2~24960784 | Chr2 | 24960784 | NA | NA | 6.21E-05 | 0.661 | NA | NA | NA | NA | Upstream | NA |
| 2~35350740 | Chr2 | 35350740 | NA | NA | 2.11E-05 | 0.661 | NA | NA | NA | NA | Intergenic | NA |
| IG5~14354843 | IG5 | 14354843 | NA | NA | NA | NA | NA | NA | 1.69E-10 | 9.26E-06 | NA | *GW5 (LOC_Os05g09520)* |
| 12~7132122 | Chr12 | 7132122 | NA | NA | NA | NA | NA | NA | 7.51E-09 | 0.0002 | Intergenic | *OsCML28 (LOC_Os12g12730); CAMK-like-46 encoding gene (LOC_Os12g12860)* |
| 10~6148264 | Chr10 | 6148264 | NA | NA | NA | NA | NA | NA | 9.08E-08 | 0.00145 | Synonymous | *Ubiquitin conjugating enzyme encoding gene* (LOC_Os10g11260) |

| SNP-ID | Chr/Pseudo -chr + | Physical position (bp) | Grain length | | Grain width | | Length-to-width ratio | | Thousand-grain weight | | Structural annotation* | Candidate genes |
|---|---|---|---|---|---|---|---|---|---|---|---|---|
| | | | P-value | FDR adjusted P-values | P-value | FDR adjusted P-values | P-value | FDR adjusted P-values | P-value | FDR adjusted P-values | | |
| IG5~13672041 | IG5 | 13672041 | NA | NA | NA | NA | NA | NA | 1.06E-07 | 0.0014 | NA | NA |
| 9~12332184 | Chr 9 | 12332184 | NA | NA | NA | NA | NA | NA | 7.39E-07 | 0.008 | Intergenic | NA |
| 5~5390763 | Chr5 | 5390763 | NA | NA | NA | NA | NA | NA | 9.78E-07 | 0.0089 | Downstream | *GW5 (LOC_Os05g09520)* |
| IG5~16943583 | IG5 | 16943583 | NA | NA | NA | NA | NA | NA | 2.23E-05 | 0.1524 | NA | NA |
| 6~24535560 | Chr6 | 24535560 | NA | NA | NA | NA | NA | NA | 3.36E-05 | 0.2042 | Intergenic | NA |
| 3~7225647 | Chr3 | 7225647 | NA | NA | NA | NA | NA | NA | 4.14E-05 | 0.2268 | Intergenic | NA |
| 6~23900256 | Chr6 | 23900256 | NA | NA | NA | NA | NA | NA | 5.42E-05 | 0.2497 | Upstream | NA |
| 10~19634914 | Chr10 | 19634914 | NA | NA | NA | NA | NA | NA | 5.48E-05 | 0.249795 563 | Intergenic | NA |
| IG3~5847364 | IG3 | 5847364 | NA | NA | NA | NA | NA | NA | 6.52E-05 | 0.274304 155 | NA | NA |
| IG5~12291400 | IG5 | 12291400 | NA | NA | NA | NA | NA | NA | 8.49E-05 | 0.323929 122 | NA | NA |
| 2~25991206 | 2 | 25991206 | NA | NA | NA | NA | 2E-07 | 0.01095 | NA | NA | Intergenic | NA |
| 7~23605893 | 7 | 23605893 | NA | NA | NA | NA | 1.9E-06 | 0.05168 | NA | NA | Upstream | NA |
| unaln_IG1~478378 73 | 16 | 47837873 | NA | NA | NA | NA | 3.8E-06 | 0.06959 | NA | NA | NA | NA |
| unaln_AROG11~1 1935468 | 14 | 11935468 | NA | NA | NA | NA | 7.9E-06 | 0.10769 | NA | NA | NA | NA |
| unaln_IG5~143548 43 | 20 | 14354843 | NA | NA | NA | NA | 3E-05 | 0.32688 | NA | NA | NA | NA |

Chr: Chromosome; Pseudo-chr: Pseudo-chromosome; 5′ UTR/3′ UTR: five prime untranslated region/ three prime untranslated region defined as per MSU7.0 annotation; Upstream: SNPs located >2 kb upstream from transcription start site of a gene; Downstream: SNPs located >2 kb downstream from transcription stop site of a gene; IG: Indica Group; JG: Japonica Group; Adm: Admixture *Structural annotation performed using MSU 7.0; #Rice Annotation Project locus ID; The loci detected from pseudo-chromosomes are highlighted in red.
