## Supplemental Table 10 for "Rice Pan-genome Array (RPGA): an efficient genotyping solution for pan-genome-based accelerated crop improvement in rice"

**Supplementary Table 10.** Details of QTLs identified for grain weight and grain size traits utilizing Sonasal × Pusa Basmati 1121 RILs

| **Trait** | **QTL** | **Chr** | **Start-End position (cM)** | **Genetic Interval (cM)** | **Start-End position (bp)** | **Physical Interval (bp)** | **LOD** | **PVE** | **Add** | **Number of genes** | **Candidate genes** |
|---|---|---|---|---|---|---|---|---|---|---|---|
| Thousand Grain Weight | *qTGW3* | 3 | 68.24-70.14 | 1.9 | 16056008-16760451 | 704443 | 13.2 | 35.7 | 2.79 | 94 | *Grain Size 3* |
| Grain Length | *qGL3* | 3 | 68.24-70.14 | 1.9 | 16056008-16760451 | 704443 | 18.9 | 42.0 | 1.04 | 94 | *Grain Size 3* |
| Grain Length | *qGL7* | 7 | 61.68-67.38 | 5.7 | 24381212-25779322 | 1398110 | 3.3 | 9.7 | 0.49 | 218 | Slender LG7 |
| Grain Width | *qGW7* | 7 | 49.61-61.68 | 6.8 | 19854664-24381212 | 4526548 | 4.9 | 13.3 | -0.08 | 708 | *OsBZR1*; *OsGRS9* |
| Grain Length to Width Ratio | *qLWR3* | 3 | 67.87-67.11 | 0.76 | 17120362-17948581 | 828219 | 15.5 | 33.4 | 0.47 | 129 | NA |
| Grain Length to Width Ratio | *qLWR7* | 7 | 62.04-62.39 | 0.35 | 24405861-24591612 | 185751 | 5.4 | 9.8 | 0.26 | 31 | NA |
