## Supplementary Results for "Rice Pan-genome Array (RPGA): an efficient genotyping solution for pan-genome-based accelerated crop improvement in rice"

### Large-scale validation of RPGA

To conduct large-scale validation of RPGA, genotyping of 188 RILs (Sonasal × Pusa Basmati 1121 $F_{12}$) and diversity panel consisting of 275 diverse rice accessions were performed using RPGA. During subsequent quality control, three RIL samples did not pass the dish quality control check (DQC < 0.82), whereas eleven RIL samples showed a lower call rate (< 97%) and thus, were eliminated from further analysis. Out of 275 natural rice accessions, only one sample failed to pass the dish quality control (DQC < 0.82) threshold whereas, only three samples failed to pass the QC call rate threshold (< 97%). A closer evaluation revealed that all the samples failing sample QC had lower quality of initial samples. The total of 177 RILs and 272 accessions from the diversity panel passed the initial quality control. Further, for both RILs and diversity panel, 80504 SNPs assayed were distinctly classified into six different classes [Poly High Resolution (PHR), Mono-High-Resolution (MHR), No Minor Homozygotes (NMH), Call Rate Below Threshold (CRBT), Off Target Variants (OTVs) and Others] according to their clustering patterns (**Figure 2**). In the case of both RILs and diversity panel, very high proportion of SNPs i.e. ~ 57% (46523) and ~ 60% (48133) of total 80504 SNPs, respectively, were classified as PHR (**Supplementary Table 1, 2**). The PHR SNPs represent the high-quality polymorphic SNPs with well-resolved clusters and, therefore, only class of SNPs usually recommended for downstream analysis. However, recently another class of SNPs called OTVs, characterized by highly reproducible but atypical pattern of reduced intensities caused by variation (PAVs/CNVs) within or distal regions of probe hybridization site, was successfully genotyped and utilized for downstream analysis (Didion et al., 2012; Mabire et al., 2019). About 25% of probes on RPGA target sub-population specific sequences from 3K rice pan-genome are, therefore, most likely to harbor PAVs/CNVs and thus expected to be classified as OTVs (**Supplementary Table 1, 2**). In the case of the diversity panel, ~15% (12618) of total 80504 SNPs were classified as OTVs whereas in the case of RILs, ~7% of total 80504 SNPs (5746) were classified as OTVs. These OTVs were then genotyped using the OTV caller provided as part of Axiom Analysis Suite to obtain genotype calls (presence/absence). Thus, 60751 (75.46%) and 52269 (65.68%) of total 80504 SNPs from the diversity panel (275 accessions) and RILs (188 individuals) genotype datasets, respectively, were selected for the downstream analysis (**Supplementary Table 1, 2**). These results confirm the high polymorphism potential and genotyping success rate of RPGA for both bi-parental mapping populations and diverse natural accessions.

Further, the genotyping error rate of RPGA-based genotyping data was assessed by comparing genotype calls of the PHR and MHR SNPs with the corresponding genotype calls generated using high-coverage whole-genome sequencing data of five accessions (name of accessions) including two parental accessions from the mapping population. This analysis revealed > 98.5% concordance for both SNPs and SVs between the two datasets. Similarly, technical replicates processed independent batches also revealed high concordance (> 99%). Thus, RPGA-based genotyping was found to have an extremely low genotyping error rate and inter-batch variation and, therefore, can be reliably used for large-scale genotyping in rice.

**Validation of RPGA-based ultra-high-density genetic map using *de novo* genome assembly of "Sonasal"**

To validate the accuracy of the aforementioned Sonsal × PB 1121 ultra-high-density genetic linkage map, the *de novo* genome assembly for one of the parental accession, Sonasal, was generated using Oxford Nanopore long-read sequencing. For this, a total of 2793046 sequence reads with a median read length > 14 kb, amounting to 48.3 Gb of raw data were generated. The N50 of sequence data was found to be > 19 kb whereas the mean read quality was estimated to be ~ 9.2 kb. The high-quality reads for assembly, only 2100126 (~ 42 Gb) reads which passed the quality filtering criterion (read length > 5 kb and qscore ≥ 8) were utilized, resulting in genome coverage of ~ 97 X. The filtered reads were then corrected based on consensus and assembled to obtain initial assembly. After initial genome assembly and subsequent polishing with Nanopore long reads as well as Illumina short reads, obtained a polished Sonasal genome assembly spanning 368.2 Mb with 304 contigs. The genome assembly displayed an N50 of 4.02 Mb indicating high contiguity. Benchmarking Universal Single-Copy Orthologs (BUSCO) analysis (Simão et al., 2015) of the Sonasal genome assembly revealed recovery of 96.5% of the 1614 BUSCOs belonging to the embryophyte group. These BUSCO statistics are comparable to previously assembled rice genomes including *japonica* cultivar Nipponbare (98.4%), *indica* cultivar R498 (98.0%), Basmati cultivar Basmati 334 (97%) and *Sadri* cultivar Dom Sufid (97%) (Choi et al., 2020). The previously developed Sonsal × PB 1121 ultra-high-density genetic linkage map was further utilized for the anchoring of contigs. From 368.2 Mb of total contig sequences, the high-density genetic map-based anchoring assigned ~ 343 Mb (~ 93%) of contig sequences to twelve linkage groups corresponding to twelve rice chromosomes. The minimal conflict was observed between marker order obtained using genetic map and marker order within *de novo* assembled contigs of Sonasal genome with < 0.2 % of makers from the genetic map not agreeing with order within contigs. This confirms the high accuracy of the previously constituted RPGA-based ultra-high-density genetic linkage map and thus, provides reliable estimates of physical locations of sub-population specific contigs from 3K rice pan-genome. This is especially crucial for the identification of dispensable genes regulating important traits using forward genetics approaches like QTL mapping and GWAS.

The chromosome level Sonasal assembly was further annotated based on available rice gene models/transcripts/proteins, utilizing the genome annotation pipeline (Holt & Yandell, 2011). This analysis identified a total of 41020 genes in the Sonasal genome. The BUSCO gene completion analysis showed that the identified genes from the Sonasal genome represent 95.2% of the 3278 single-copy genes from the Liliopsida gene dataset. Further, to assess genomic similarities and the syntenic relationship between Sonasal and other available genomes of rice accessions from the diverse varietal groups, the Sonasal draft genome was compared to the Nipponbare (*japonica*), Nagina 22 (*aus*), IR 64 (*indica*), Basmati 334 and Dom Sufid (aromatic) reference genomes using whole-genome alignment. The comparison revealed a high degree of microsynteny between the Sonasal draft genome and the Nipponbare genome (**Figure 4**). These further highlights the utility of RPGA-based ultra-high density genetic linkage maps coupled with long-read sequencing, for developing high-quality chromosome-level genome assemblies.

#### Nipponbare-specific gene loci detected using pan-genome-based GWAS

#### ***OsGAMYB1*** associated with rice grain length

A grain length-associated locus containing most significant SNP (MSP) was detected on chromosome 1 (MSP-Chr1: 39294258). The locus was found to harbor 29 protein-coding genes. Among these genes, *OsGAMYB1*, a gene encoding MYB transcription factor (*LOC_Os01g59660*) was found to be located 26.22 kb upstream from MSP-Chr1: 39294258. The *in silico* expression analysis revealed that *OsGAMYB1* predominantly expresses during panicle and seed development. MYB transcription factors are well known to regulate seed size in *Arabidopsis* (Zhang et al., 2013c). Further, *miR159* is known to regulate seed size along with flowering time, anther development and vegetative growth by targeting various *MYB* genes (*AtMYB56* and *AtMYB65*) in *Arabidopsis* (Allen et al., 2007; Palatnik et al., 2007). Interestingly, this *miR159-MYB* module was found to be conserved in rice, where, *OsmiR159* has been shown to target *OsGAMYB1* (*LOC_Os01g59660*) which is orthologous to *AtMYB65*, and another *MYB* gene *OsGAMYB-like1* (*LOC_Os06g40330*). The suppression of *miR159* has been shown to reduce grain length (12.68–16.90%), grain width (6.06–9.09%) and grain thickness (15.51–16.35%), ultimately leading to a reduction in thousand-grain weight (32.11–33.58%) (Zhao et al., 2017). This evidence makes *OsGAMYB1* a likely candidate gene regulating grain size in rice. Further, fine mapping of this locus will be required to authenticate *OsGAMYB1* (*LOC_Os01g59660*) as a major effect gene and to identify the causal mutation regulating grain size/weight in rice.

#### Serine/threonine-protein kinase associated with rice grain width

Similar to grain length, prominent grain width associated locus on chromosome 11 was found to harbor serine/threonine-protein kinase gene (*LOC_Os11g40970*). Previously, genes encoding diverse serine/threonine-protein kinases are reported to regulate seed development in both *Arabidopsis* and rice, indicating their important role in seed development both in monocots and dicots (Duan et al., 2014; Xu et al., 2018a,b; Chun et al., 2020; Zhang et al., 2020). Interestingly, *LOC_Os11g40970* was found to be one of the genes down-regulated in *gibberellin-deficient dwarf1 (gdd1)* mutant that has reduced root length, stem and smaller seeds, compared to wild type due to reduced cell size (Li et al., 2011a). Further, *LOC_Os11g40970* expresses predominantly in leaf, panicle and spikelet. Therefore, *LOC_Os11g40970* can be a probable candidate gene regulating grain width in rice.

**F-box and *OsRUB1* associated with rice grain weight**

Among loci detected for thousand-grain weight, a prominent locus on chromosome 4 (MSP-Chr4:4486000) that harbored a total of 17 protein-coding genes was investigated further. These genes included two probable F-box protein-encoding genes. One of these F-box genes (*LOC_Os04g08460*) displays higher expression during panicle and spikelet development. F-box proteins are essential components of the ubiquitin-proteasome pathway, a major pathway known to regulate grain size/weight in rice (Song et al. 2007; Li et al., 2012; Huang et al., 2017; Choi et al., 2018; Shi et al., 2019). Therefore, it would be interesting to further investigate the role of *LOC_Os04g08460* in grain size/weight regulation in rice.

Further, the thousand-grain weight associated locus on chromosome 10 (MSP-Chr10:21588761) harbored 13 protein-coding genes. Among these genes, *LOC_Os10g11260* was found to encode ubiquitin-conjugating enzyme, an ortholog of *Arabidopsis RELATED-TO-UBIQUITIN* (*RUB1*). *RUB1* encodes an ubiquitin-like protein that acts as a vital component of the SCF complex (ubiquitin ligase) and it is known to be essential for early embryonic cell division during seed development (Bostick et al., 2004). This suggests the probable role of *RUB1* (LOC_Os04g08460) in regulating the thousand-grain weight in rice.

**Novel sub-population specific (dispensable) genes regulating rice grain size/weight identified using RPGA-based GWAS**

***WDR12* and *GRAS* associated with rice grain length**

For rice grain length, two important loci (MSP-unaln_JG10~4245875 and MSP unaln_JG10~6950498) were detected on the pseudo-chromosome JG10 (**Figure 7; Supplementary Figure 4, 5, 6, 7; Supplementary Table 5, 6, 7, 8, 9**). The physical position of these SNP loci was further determined with reference to the Nipponbare genome using an integrated genomic approach (Detailed strategy as per Materials and methods). Based on these analyses, WD repeat-containing PROTEIN 12 (*LOC_Os07g40930*) and GRAS family transcription factor (*OsR498G0612839600*.01) are the most likely genes for MSP-unaln_JG10~4245875 and MSP unaln_JG10~6950498 loci, respectively governing grain length in rice.

**Multi-domain protein associated with rice grain width**

Similar to grain length, two SNPs (MSP-unaln_IG2~18885542 and MSP-unaln_IG2~35461875) from subpopulation-specific pseudo-chromosome IG2 (*indica* group 2) displayed significant association with grain width (**Figure 7; Supplementary Figure 4, 5, 6, 7; Supplementary Table 5, 6, 7, 8, 9**). One of these associations which crossed the stringent significance threshold (MSP-unaln_IG2~18885542) was further investigated to determine probable candidate gene. The physical positions of the contigs harboring SNP, unaln_IG2~18885542 were identified on the chromosome. The identified positions were cross-validated using information from the Sonasal × PB 1121 genetic map. The location of contig harboring SNP unaln_IG2~18885542 was found to be located on chromosome 10 and also found to overlap with a gene (*LOC_Os10g31770*) encoding a multi-domain (PH, START and DUF1136) protein. The multiple alignment of sequences from seven rice accessions, Nipponbare (*japonica*), *indica* (IR 64 and R 498), *aus* (Nagina 22) and aromatic (Sonasal, Basmati 334, and Dom Sufid) revealed the presence of multiple InDels and SNPs within the intronic, 3' UTR and downstream region of this region. Further analysis suggested that the aforementioned sequence variation leads to loss of PH domain in all four compared accessions (Name), except Nipponbare. The PH domain-containing proteins had been previously reported to function in cellular signal transduction pathways by interacting with proteins such as βγ-subunits of heterotrimeric G proteins and protein kinase C. Various G protein subunits are already been reported to play important role in the regulation of grain size/weight in rice. In addition to this, *LOC_Os10g31770* was shown to act downstream of *RICE OUTERMOST CELL-SPECIFIC GENE 5* (*ROC5*). *ROC5* regulates cell size and cell number in diverse plant organs including leaf and spikelets of rice (Zou et al., 2011). Further, *ROC5* has been believed to act downstream of *MINI SEED 2 (MIS2)*, a known gene-regulating grain size/weight in rice (Chun et al., 2020). Interestingly, 742 bp deletion at 5'end of *LOC_Os10g31770* leading to loss of complete 5'UTR was found in all four accessions compared to Nipponbare. This evidence suggests *LOC_Os10g31770* as a possible candidate gene for QTL associated with SNP-unaln_IG2~18885542.

***WDR12* and Multi-domain protein associated with rice grain length-to-width ratio**

In case of grain length-to-width ratio, apart from the aforementioned loci from the Nipponbare reference genome, three loci from sub-population specific pseudo-chromosomes were also found to be associated with length-to-width ratio (**Figure 7; Supplementary Figure 4, 5, 6, 7; Supplementary Table 5, 6, 7, 8, 9**). Among these, two loci (MSP-unaln_JG10~6950498 and MSP-unaln_IG2~18885542) were repeatedly detected with different statistical models of GWAS.

Interestingly, these two loci i.e., SNP-unaln_JG10~6950498 and SNP-unaln_IG2~18885542, were also detected for grain length and grain width traits, respectively. The WD repeat-containing PROTEIN 12 (*LOC_Os07g40930*) encoding gene and a gene (*LOC_Os10g31770*) encoding a multi-domain (PH, START and DUF1136) protein were identified as most likely candidates for MSP-unaln_JG10~6950498 and MSP-unaln_IG2~18885542, respectively. As previously discussed, WDR12 proteins are an essential part of the PeBoW (PES-BOP1-WDR12) complex. The reduction in PeBoW proteins was found to suppress cell proliferation and cell expansion in *Arabidopsis* underlining its vital role in the regulation of cell cycle in plants (Cho et al., 2013; Ahn et al., 2016). Thus, it will be interesting to further evaluate the function of *LOC_Os07g40930* (WDR12 protein-encoding gene) and *LOC_Os10g31770* (multi-domain protein-encoding genes) in regulating grain size (grain length/length-to-width ratio) of rice.

**Dirigent and unknown expressed genes associated with rice thousand-grain weight**

For thousand-grain weight, two significant associations (SNP-unaln_IG5~14354843 and SNP-unaln_JG10~6243872) were detected from two different subpopulation specific pseudo-chromosomes i.e. IG5 and JG10 (**Figure 7; Supplementary Figure 4, 5, 6, 7; Supplementary Table 5, 6, 7, 8, 9)**. As the chromosome of origin and physical locations of these SNPs are unknown, efforts were made to ascertain the physical locations of these SNPs (QTLs thereof) with reference to the Nipponbare reference genome. For this, the pair-wise LD statistics, BLAST and multiple sequence analysis were utilized as previously discussed. The analysis suggested that the location of SNP-unaln_IG5~14354843 coincides with *GW5* locus which has also been detected with SNPs from the Nipponbare reference genome. The 1212 bp InDel (SV) located upstream of *GW5* (calmodulin binding protein-encoding gene) has been identified as a causal variation responsible for regulating grain size (Liu et al., 2017). This further proves the efficacy of the RPGA-based GWAS approach used in current study to efficiently identify SVs associated with important agronomic traits. Further, the second SNP, unaln_JG10~6243872 was found to be located on chromosome 12. The contig containing SNP-unaln_JG10~6243872 spans the intergenic region between a dirigent family gene (*LOC_Os12g12600*) and the gene encoding protein of unknown function (*LOC_Os12g12610*). Although no functionally relevant variation was observed within the region spanned by contig sequence harboring SNP, unaln_JG10~6243872, a 2120 bp deletion was detected in dirigent gene (*LOC_Os12g12600*). Dirigent family genes are crucial for lignan and lignin biosynthesis and hence play an important role in lignin deposition in the cell walls (Paniagua et al., 2017). Therefore, it will be interesting to study the potential role of the dirigent gene in the regulation of grain weight in rice.
