## Supplementary Method for "Rice Pan-genome Array (RPGA): an efficient genotyping solution for pan-genome-based accelerated crop improvement in rice"

### SUPPLEMENTARY MATERIAL AND METHODS

#### Designing of a rice pan-genome array (RPGA) for SNP genotyping

About 30 M SNPs common to all aforementioned SNP datasets were selected and filtered based on the following criteria. (I) considered only biallelic SNPs, (II) eliminated SNPs with > 30% missing data, (III) romoved SNPs with minor allele frequency less than 10% (MAF < 0.1), (IV) 25-bp sequence flanking the interrogated SNPs should be invariant (at least on one side of interrogated SNP) and should have 40 to 70% GC content, and (V) the 25-bp flanking sequences of interrogated SNPs should not have any other genomic match ≥ 85% in rice pan-genome sequence. Further, the utility of these SNPs were assessed based on p-convert scores generated using the Affymetrix Power Tool (APT) AxiomGTv1 algorithm. The p-convert score represents the probability of SNPs being converted into a successful Axiom assay. For calculation of p-convert score, a random forest model is used, which takes into account various factors including sequence complexity of probes, the probability for non-specific hybridization, and binding energy, etc. Based on p-convert scores of SNPs (probes), these were classified as recommended (p-convert score > 0.6), neutral (0.6-0.4) and not-recommended (p-convert score < 0.4). All the SNPs (probes) for which both strands did not pass the p-convert score threshold of 0.4 were discarded from further analysis. The remaining SNPs were utilized for genome-wide tag SNP selection. The LD-based informative tag SNPs ($r^2 \geq 0.64$ and MAF > 0.1) were selected using LRTag which employs the Lagrangian Relaxation-based algorithm to address the minimum common tagSNP selection (MCTS) problem (Liu et al., 2007). Further, the rice 3K pan-genome sequence was divided into 13341 number of 50 kb bins and informative SNPs were further added (from any one of the aforementioned datasets) to the bins with no SNPs to ensure the maximum genome coverage with the uniform genome-wide distribution. About 2000 SNPs from 164 functionally characterized known genes (includes 2 kb each of upstream and downstream regions from the translation initiation codons of genes) associated with grain size/weight, grain quality and grain aroma traits in rice were also included. Adhering to the aforementioned criteria, 176128 probes corresponding to 80502 SNPs were selected finally for tiling on the RPGA. In addition, 3996 probes (1332 controls tiled 3 times) corresponding to 3000 monomorphic SNPs were also included as dQC monomorphic controls in the RPGA **(Supplementary Figure 13**).

#### RPGA genotyping

For genotyping with rice 80K RPGA, high-quality genomic DNA from young leaves of 190 Sonsal × Pusa Basmati 1121 RILs, two RIL parental accessions, 271 accessions from a diversity panel, five $F_1$ hybrids and three back-cross lines was extracted with DNeasy plant mini kit (Qiagen, USA) as per manufacturer's instructions. The genomic DNA isolated from samples was then quantified with a NanoDrop1000 spectrophotometer (Thermo Scientific, USA) as well as a Qubit fluorometer (Thermo Scientific, USA) and diluted to a concentration of 10 ng/ul. The 20 µl of diluted DNA was then utilized for further target preparation. Target preparation was done following Axiom manual target preparation protocol in five distinct stages, I) Whole-genome amplification of sample genomic DNA, II) Fragmentation of amplified DNA (25-125 bp fragments) and further purification using isopropanol precipitation, III) Drying and resuspension of DNA pallets (quality control with agarose gel electrophoresis for ensuring desired fragment size, and IV) Denaturation of DNA samples and preparation for hybridization. Finally, all the array processing steps including loading of hybridization ready samples on to an 80K RPGA, ligation, washing, staining (with two different dyes) and subsequent scanning of the array to detect hybridization signals were performed using GeneTitan® Multi Channel (MC) instrument (Affymetrix, USA) following manufacturer's instructions. The CEL files (for all the genotyped) generated after processing of five set of RPGAs in the GeneTitan were further used for obtaining genotype calls.

**RPGA data analysis and genotype data imputation**

For genotype calling, CEL files generated previously were uploaded to Axiom Analysis Suite 2.0 (Affymetrix, USA) and further analysis was performed following the Axiom best practices workflow. Initially, high-quality samples were filtered based on the DQC and QC call rate. DQC (Dish-QC) values represent the resolution of "contrasts" values i.e. the difference in signal intensities between two types of non-polymorphic probes which includes AT non-polymorphic probes and GC non-polymorphic probes. DQC value ranges between zero and one, where zero represents no resolution and 1 represents complete resolution, between distributions of AT and GC probe contrast values. However, DQC is often not sufficient to detect all low-quality samples, therefore further sample filtering was done based on QC call rate [number of samples assigned with genotype call (AA, AB, BB) divided by the total number of samples] of a subset of probes with well-resolved clusters. For both DQC and QC call rates, filtering default values of >0.82 and ≥97, respectively, were utilized. All the samples failing any of these filterings were eliminated from further analysis. Genotype calls for all the QC passed samples were then produced using the AxiomGT1 algorithm utilizing the generic priors i.e., the same prepositioned cluster location is provided to clustering algorithm for computation of three posterior cluster locations for each SNP.

Further, SNP QC was performed on available genotype data utilizing *Ps_Metrics* and *Ps_Classification* functions available as part of Axiom Analysis Suite 2.0 (Affymetrix, USA). *Ps_Metrics* function computes 12 different SNP matrices for each probe set genotyped in the previous step. Out of these 12 SNP matrices, four [Call Rate (CR), Fisher's Linear Discriminant (FLD), Heterozygous Strength Offset (HetSO), Homozygous Ratio Offset (HomRO)] are utilized by *Ps_Classification* function for classification of each probe-set into one of the six different categories. For SNP QC, default parameters recommended for diploid species (except HetSO, which was set to ≥ -0.3 instead of default ≥ -0.1) were utilized. The parameters include SNP call rate (CR) cutoff ≥ 97, Fisher's Linear Discriminant (FLD) cutoff ≥ 3.6, Heterozygous Strength Offset (HetSO) cutoff ≥ -0.3, Heterozygous Strength Offset Off Target Variant (HetSO.OTV) cutoff ≥ -0.3, Homozygote Ratio Offset 1 (HomRO-1) cutoff ≥ 0.6, Homozygote Ratio Offset 2 (HomRo) cutoff ≥ 0.3, and Homozygote Ratio Offset (Homo RO) 3 cutoff ≥ -0.9. This includes Poly High Resolution (well-resolved clusters with at least two observation of minor alleles), No Minor Homozygous (well-resolved clusters with no observation of homozygous minor allele), Off Target Variants (middle AB cluster is split into additional cluster i.e. OTV cluster, other than three usual clusters), Mono High Resolution (presence of single cluster representing one of the two alleles), Call Rate Below Threshold (well-resolved clusters, however, call rate is below threshold) and other (one or more cluster properties are below threshold values). In addition, an inbred penalty of 4 was applied to all the samples considering their highly homozygous nature (except for backcross lines and $F_1$ hybrids). Further, SNP cluster plots were visualized employing the *Ps_Visualization* function integrated into Axiom Analysis Suite 2.0. Before proceeding further, a few SNPs (~200) from all the classes were visually inspected to ensure optimal classification by standard SNP QC filters. Further, few additional sample QC steps (samples with unusually high relatedness and displaying high DQC but low QC call rates were detected as contaminated) as well as plate QC steps (based on DQC distribution and plate-wise MAF differences to detect problems during processing of plates) were performed. After selecting high-quality plate/samples, SNPs belonging to recommended categories for diploid, such as Poly High Resolution, Mono High Resolution and Off-Target Variants (after genotyping with OTV caller function) were further selected for downstream analysis. The genotype calls of RPGA for passing samples and recommended SNPs were then exported in .TXT format.

**High-density genetic linkage map construction**

To construct a high-density genetic linkage map, 80K RPGA-derived genotype data of 190 (PB 1121 × Sonasal) RILs was filtered to keep only polymorphic SNPs between two parental accessions converted to ABH genotype format using TASSEL 5.0 (Bradbury et al., 2007). The ABH formatted genotype data was then used as input to the R/qtl program (Broman et al., 2003) for the genetic map construction. Before initiating map construction, all the SNPs and RILs with > 10% missing genotyping data were eliminated. Further, RILs displaying a high level of genetic similarity (>95%) were omitted from analysis. In addition, SNPs displaying significant deviation from the expected 1:1 segregation ratio for RILs ($P$ > 0.01) were identified with the *chi-squared* test and removed from further analysis. After initial quality control of genotype data, pair-wise recombination fraction (RF) and the logarithm of odds (LOD) scores were calculated for all SNP marker-pairs. Linkage groups were then formed by grouping marker-pairs with the minimum LOD of 8 and maximum RF of 0.40. Markers from individual linkage groups were then ordered to obtain a preliminary genetic map. This preliminary genetic map was then utilized for the detection of problematic RILs (RILs with abnormally high/low crossovers) and potential genotype errors markers. These problematic RILs were eliminated from further analysis. Detected genotype errors were also eliminated by setting all potential erroneous genotypes to missing. The re-ordering of markers and rippling were performed with a window size of 5 to ensure optimal marker order and finally a RPGA-based high-density genetic linkage map was constructed.

**High molecular DNA isolation for Nanopore sequencing**

The 400 mg of leaf tissue from twenty different 25-days-old plants of a short-grain rice variety Sonasal were harvested, pooled, flash-frozen using liquid Nitrogen and stored at -80 $^{0}$C. The DNA extraction was carried out by nuclei isolation, lysis and column purification (https://stories.rbge.org.uk/archives/30792). The quality as well as quantity of DNA isolated was assessed using NanoDrop 1000 Spectrophotometer (Thermo Fisher Scientific, USA). Quantification of extracted genomic DNA was further performed using the Qubit 2.0 (Thermo Fisher Scientific, USA) fluorometer. Finally, the integrity of genomic DNA was assessed by agarose gel electrophoresis.

**Long-read Nanopore and short-read Illumina sequencing**

For Nanopore sequencing, the high-quality DNA isolated from Sonasal was used for constructing genomic DNA library using the SQK-LSK109 ligation kit. The libraries were further sequenced using FLO-MIN106 (R9.4) flowcells on PromethION 24 (P24) platform. For short-read Illumina sequencing, the genomic DNA was fragmented to 300-bp size using a Covaris ultrasonicator. The libraries were prepared using Illumina TruSeq DNA sample prep kit (Illumina, USA) following the manufacturer's protocol. The size distribution of the libraries was assessed using a Bioanalyzer 2100 (Agilent Technologies, USA). The libraries were then sequenced using the Illumina HiSeq2000 platform (Illumina, USA) by paired-end sequencing.

**Genome assembly, polishing and annotation**

The raw signal intensity data (FAST5) of long-read sequences generated by Nanopore was base called with the flip flop v2.3.5 program. The quality statistics of the base called data were visualized with nanoQC (De Coster et al., 2018). Further, reads with a quality score < 8 and read length < 5 kb were filtered out using NanoFilt nanoQC (De Coster et al., 2018). The reads remaining after filtration was then corrected using Canu (Koren et al., 2014). The corrected reads were then assembled using Flye genome assembler (Kolmogorov et al., 2019). The assemblies produced were then polished with Recon v1.4.11, for three rounds and with Medaka (https://github.com/nanoporetech/medaka), for one round.

The Illumina short-reads were then mapped onto the draft assemblies using bwa-mem (Li and Durbin, 2009). The alignment file generated was then used by Pilon v1.22 (Walker et al., 2014), for three rounds of final polishing. The calculation of genome assembly statistics was performed with help of bbmap stats.sh script available as a part of BBTools (https://jgi.doe.gov/data-and-tools/bbtools/). BUSCO v4.0.5 (Simão et al., 2015) was used to assess the completeness of genome assemblies. D-GENIES program (Cabanettes and Klopp, 2018) was used to analyze the synteny between the Sonasal draft genome and the Nipponabare genome (IRGSP 1.0). The Sonasal draft genome was annotated utilizing a MAKER annotation pipeline (Holt and Yandell, 2011) to identify protein coding sequences present in the Sonasal genome. The publicly available rice cDNA sequences (https://rapdb.dna.affrc.go.jp/) were supplied as the EST evidence and the protein sequences from 13 *Oryza* species were provided as the protein evidence (Stein et al., 2018). RepeatMasker v4.0.7 was used to assess the repeat content based on previously developed *de novo* repeat libraries for different wild and cultivated members of *Oryza species* (Choi et al., 2017). The repetitive regions were then masked out before further gene prediction. The genes predicted in the first round of MAKER were subsequently used as a training dataset for SNAP (Korf, 2004) and Augustus (Stanke et al., 2008) during the second round of gene annotation with MAKER. The high-density genetic linkage maps developed from Sonasal × PB 1121 RIL mapping population were then used for the scaffolding of annotated contigs with Chromonomer v1.10 (Catchen et al., 2020).

**Population structure and molecular diversity analysis**

To decipher the population structure in 271 diverse Indian accessions, a core SNP dataset was generated with the LD pruning procedure implemented in PLINK v2.0 (Chang et al., 2015). During this procedure, a window size of 10 kb, window step of 1 SNP and $r^2$ threshold of 0.8 was employed to keep only independent SNPs. This core SNP dataset (5812 SNPs) was further utilized for multi-dimensional scaling (MDS) and population structure analysis. For MDS analysis, IBS (identity by state) distance matrix was calculated with PLINK v2.0 (Chang et al., 2015) which was further used to perform MDS analysis using 'cmdscale' function form 'stats v.3.6.2' R package. To determine population structure, the core SNP dataset was analyzed utilizing the variational Bayesian model of fastSTRUCTURE v1.0. The analysis was performed with default fastSTRUCTURE v1.0 parameters for convergence criterion and priors for the number of clusters (K) ranging from 1 to 15. The most-likely K value was determined based on the best chose function of fastSTRUCTURE. Finally, the replicates of chosen clusters were summarized with the CLUMPK program (http://clumpak.tau.ac.il/).

**Identification of candidate genes underlying trait associated genomic loci**

To select candidate genes underlying associated loci from subpopulation-specific pseudo-chromosomes, the Nipponbare reference genome-based method for candidate gene selection could not be utilized. This is due to absence of these associated loci from the Nipponbare reference genome and unknown physical position of these associations relative to Nipponbare genome. Therefore, candidate genes underlying associated loci from subpopulation-specific pseudo-chromosomes were determined using an integrated approach based on multiple rice reference genomes. This integrated approach described here assesses the novelty of association and identifies underlying candidate genes, 1) The approximate location of MSPs relative to Nipponbare reference genome were estimated based on pair-wise LD between MSP and all SNPs belonging to Nipponbare reference genome as follows, 2) The estimated location of MSPs were then revalidated with ultra-high-density genetic linkage map developed using Sonasal × PB 1121, 3) MSPs form 12 pseudo-chromsomes colocalizing with any of the associated loci identified from the Nipponbare genome were eliminated from further analysis, as these does not represent novel association, 4) The physical positions of pseudo-chromosomes contigs harbouring the MSPs corresponding to novel association were then identified in the genome assemblies of six different accessions belonging to different rice subpopulations that include Nipponbare (*japonica*), Nagina 22 (*aus*), IR 64 (*indica*), Basmati 334, and Dom Sufid, Sonasal (aromatic) using BLAST, 5) The complete nucleotide sequence including 3 kb flanking sequences were retrieved from each of the six genome assemblies based on their respective positions, 6) The sequences retrieved form six different genome assemblies and pseudo-chromosome contigs were subjected to multiple sequence alignments, and 7) The candidate genes were then predicted based on sequence variation (especially INDELs) observed between multiple reference genomes within aligned genomic regions.
